# Convergence in form and function overcomes non-parallel evolutionary histories in a Holarctic fish

**DOI:** 10.1101/265272

**Authors:** Arne Jacobs, Madeleine Carruthers, Andrey Yurchenko, Natalia V. Gordeeva, Sergei S. Alekseyev, Oliver Hooker, Jong S. Leong, David R. Minkley, Eric B. Rondeau, Ben F. Koop, Colin E. Adams, Kathryn R. Elmer

**Affiliations:** Institute of Biodiversity, Animal Health and Comparative Medicine, University of Glasgow, G12 8QQ, Glasgow, UK; Vavilov Institute of General Genetics, Russian Academy of Sciences, ul. Gubkina 3, Moscow, 199333 Russia; Koltzov Institute of Developmental Biology, Russian Academy of Sciences, ul. Vavilova 26, Moscow, 119991 Russia; Severtsov Institute of Ecology and Evolution, Russian Academy of Sciences, Leninskii prosp. 33, Moscow, 119071 Russia; Scottish Centre for Ecology and the Natural Environment, University of Glasgow, Rowardennan, Loch Lomond, Glasgow G63 0AW, UK; Biology/Centre for Biomedical Research, University of Victoria, British Columbia, Canada; PR statistics, 6 Hope Park Crescent, Edinburgh, EH8 9NA, UK

## Abstract

Understanding the extent to which evolution is predictable under multifarious selection is a longstanding question in evolutionary biology. However, the interplay of stochastic and contingent factors influencing the extent of parallelism in nature is not well understood. To test the predictability of evolution, we studied a ‘natural experiment’ on different organismal levels across lakes and evolutionary lineages of a freshwater salmonid fish, Arctic charr *(Salvelinus alpinus).* We identified significant phenotypic parallelism between Arctic charr ecotype pairs within a continuum of parallel evolution and highly parallel adaptive morphological traits. Variability in phenotypic predictability was explained by complex demographic histories, differing genomic backgrounds and genomic responses to selection, variable genetic associations with ecotype, and environmental variation. Remarkably, gene expression was highly similar across ecotype replicates, and explained the observed parallelism continuum. Our findings suggest that parallel evolution by non-parallel evolutionary routes is possible when the regulatory molecular phenotype compensates for divergent histories.

## Background

The degree to which the pathways of evolution are predictable, particularly under complex natural conditions, remains one of the greatest questions in evolutionary biology(Conway Morris 2003; Gould 1990). Numerous species in nature have repeatedly evolved similar phenotypes in response to similar environmental challenges, strongly suggesting repeatability and predictability of evolutionary trajectories (Elmer & Meyer 2011; Kaeuffer et al. 2012; Mahler et al. 2013; Kowalko et al. 2013; Elmer et al. 2014), and highlighting the pervasive role of natural selection in evolution (Schluter 2001; Endler 1986). Despite this, extensive variation in the magnitude and direction of evolutionary trajectories has been observed in some classic ‘parallel’ populations, suggesting a major influence of contingency and stochasticity (Elmer et al. 2014; Oke et al. 2017; Bolnick et al. 2018; Stuart et al. 2017). Stochastic factors such as differences in the local environment, gene flow, and selection regimes, or contingencies such as genomic background, demographic history, and the genetic architecture of adaptive traits, can lead to departures from phenotypic parallelism and deviations at the genomic level (Elmer et al. 2014; Kowalko et al. 2013; Conte et al. 2012). To improve our ability to predict evolutionary outcomes, it is critical to understand the evolutionary routes leading to replicated ecological divergence in a range of independent systems and disentangle the impact of various contingent and stochastic factors on the degree of phenotypic parallelism.

Predictability of evolution when selection is multifarious and traits are quantitative is not well understood and is difficult to quantify in a biologically relevant way (Nosil et al. 2018; Oke et al. 2017). Multiple statistical frameworks have been developed recently to address parallel evolution in natural systems, as a proxy for predictability of evolution, consistency of selection gradients, and pervasiveness of deterministic processes (Collyer & Adams 2013; Langerhans & DeWitt 2004). Parallel evolution of phenotypes can be inferred from heritable, ecologically relevant traits, overall morphology, and/or ecological niche (Stuart et al. 2017; Elmer et al. 2014; Soria-Carrasco et al. 2014; Linnen et al. 2009) and is quantified as replicated evolutionary trajectories (Collyer & Adams 2013) or as the amount of variation explained by ecotype (Langerhans & DeWitt 2004). Parallel evolution at the genetic level is evidenced by shared genes and genetic regions underpinning parallel phenotypes across replicates, either due to repeated *de novo* mutation or recruitment of shared standing genetic variation (Elmer & Meyer 2011; Conte et al. 2012). We describe parallel evolution as the replicated, independent, evolution of quantitatively similar adaptive phenotypes; i.e. not only the outcome but also the process. The study of replicated natural systems at different organismal levels is critical to identifying the genetic, environmental, and selective components underlying parallel evolution and its deviations.

The replicated ecological and morphological post-glacial diversification of fishes into distinct trophic specialists in freshwater lakes is a powerful natural experiment for testing phenotypic and evolutionary predictability (Schluter 1996; Jonsson & Jonsson 2001; Alekseyev et al. 2002). In the Holarctic salmonid species Arctic charr *(Salvelinus alpinus)*, sympatric ecotype pairs that differ in several heritable phenotypic traits, such as body shape, body size, trophically-relevant morphology, and life history, are abundantly replicated (Jonsson & Jonsson 2001; Alekseyev et al. 2002; Adams & Huntingford 2002; Garduño-Paz et al. 2012; Hooker et al. 2016). Ecotypes distinguished by diet and foraging tactics include: Planktivores and Piscivores which feed in the pelagic, and Benthivores which feed in the benthic-profundal zone (Jonsson & Jonsson 2001; Alekseyev et al. 2002; Adams & Huntingford 2002; Garduño-Paz et al. 2012). These ecotypes can be found in several Arctic charr lineages (Jonsson & Jonsson 2001; Alekseyev et al. 2002), but are most frequent in the Atlantic and Siberian lineages, which most likely diverged before the last glacial maximum during the Pleistocene (Lecaudey et al. 2018). Arctic charr ecotypes have most likely evolved following the last glacial maximum (around 10,000 – 20,000 years ago), after the colonization of newly formed postglacial lakes through putatively pelagic charr from different glacial refugia populations. However, it is not known how phenotypically parallel these ecotypes are across lakes and lineages, if they have evolved in sympatry as a divergence of a single refugial population or following the secondary contact two or more refugial populations, and if similar genomic regions and genes are involved in the divergence.

Here we tested the extent of parallelism in phenotype, evolutionary history, genomic basis, and transcriptomic expression in repeated divergences of Arctic charr in their environmental context from eleven lakes within and across two evolutionary lineages (Atlantic and Siberian). We revealed a continuum of phenotypic parallelism and subsequently linked this to variation in the extent of parallelism in environment, evolutionary history, genomic basis and gene expression.

## Results and Discussion

### Phenotypic divergence and parallelism

To test for parallelism in ecologically relevant phenotypes, we assessed morphology based on seven linear traits for Arctic charr from Scotland and Siberia (total N=1,329 individuals). This included eleven replicate ecotype pairs: six benthivorous-planktivorous and five piscivorous-planktivorous combinations (Fig. 1; Supplementary table S1). Nine of these ecotype pairs represent independent divergences from different lakes, catchments or lineages. Additionally, three populations were included as outgroup comparisons in morphological analyses (piscivorous from Kudushkit, planktivorous ecotypes from Maloe and Bol’shoe Leprindo) because the sympatric sister ecotype is extremely rare, and one population (Tokko) with a benthivorous and mainly insectivorous ecotype that has no population replicate. These populations were excluded from the parallelism analysis but included in later genetic analyses.

**Figure 1.**
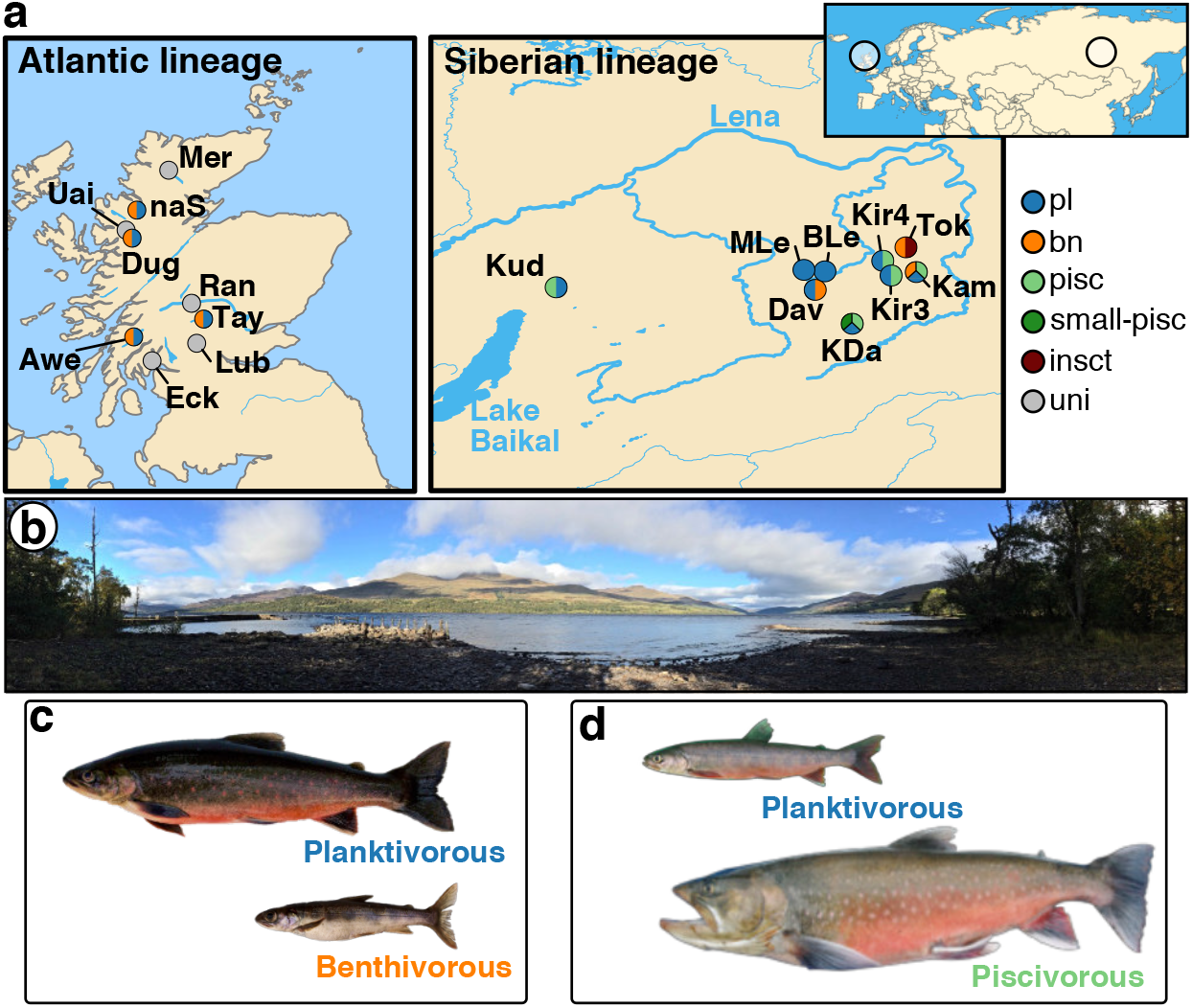
Sampling locations and ecotypes. **(a)** Maps showing the sampling locations of Arctic charr from the Atlantic lineage in Scotland (N=440) and the Siberian lineage in Transbaikalia, Russia (N=1,009). The colour combination of each dot shows the ecotypes sampled from each location. Full names of lakes are given in Supplementary table S1. **(b)** Picture showing the sampling site at Loch Tay in Scotland. (c and d) Representative individuals of the **(c)** sympatric ecotypes from Loch Tay in Scotland and the **(d)** sympatric ecotypes from Kalarskii Davatchan (KDa) in Transbaikalia.

To explore the degree of parallelism in individual traits primarily related to trophic morphology, habitat use and swimming ability, we first used a trait-by-trait linear modelling approach for all ecotype pairs. We found that ecotype (parallel divergence term) explained more phenotypic variance than the ‘ecotype x lake’ and ‘ecotype x lineage’ interaction terms (nonparallel aspects of divergence) for all traits (Fig. 2a,b, Supplementary Fig. S1a), indicating that replicated ecotypes are similar in magnitude and/or direction of divergence. The direction and magnitude of trait differences between ecotypes varied across populations, ranging from highly parallel to antiparallel in some populations and traits (Fig. 2c). Furthermore, for five of seven traits, the effect of lake (HL, ML, LJL) or lineage (HDE, HDO) had the strongest effect on trait variance (Supplementary table S2). In combination, these results suggest that trait divergence between ecotypes is predictable across lakes and lineages though the absolute trait values differ in each population. However, for eye diameter (ED) and pectoral fin length (PFL), ecotype explained the highest proportion of phenotypic variation (Fig. 2a, Supplementary Fig. S1, Supplementary table S2). This indicates that across lakes and also across evolutionary lineages ecotypes are consistently diverged in those traits, which are closely related to habitat use (ED) and swimming performance (PFL) (Klemetsen et al. 2002). This agrees with previous studies on individual Arctic charr populations and other salmonid species that found heritable, but also plastic, differences in these traits between ecotypes with profundal-benthic ecotypes having larger eyes and different pectoral fins than pelagic or littoral ecotypes, most likely as an adaptation to lower light conditions and different foraging manoeuvrability (Siwertsson et al. 2013; Klemetsen et al. 2002; Adams & Huntingford 2004).

**Figure 2.**
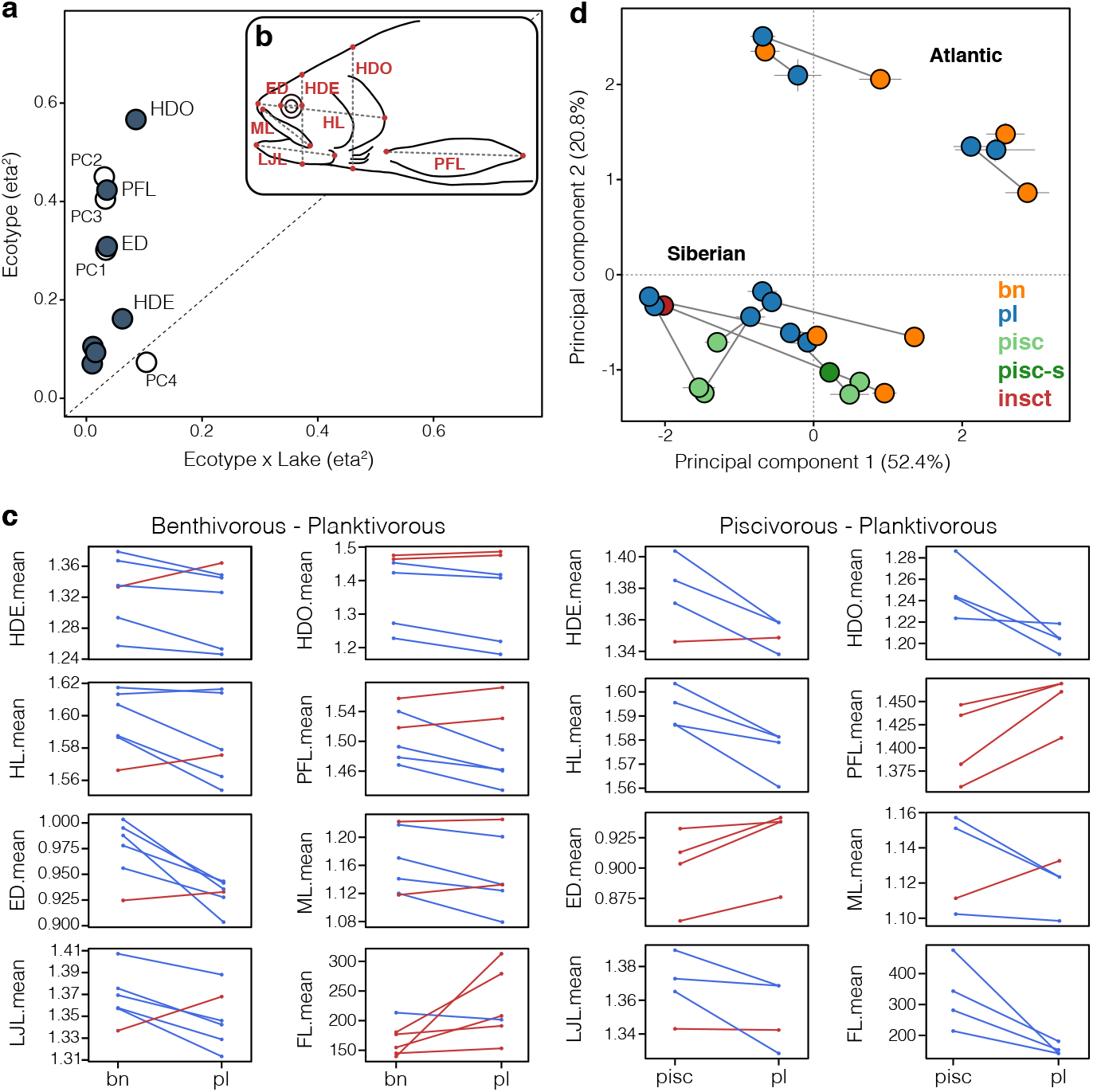
Continuum of phenotypic parallelism. **(a)** Effect sizes (partial r^2^) of the ecotype and ‘ecotype x lake’ interaction terms for all seven linear traits (dark dots) and PC1 to PC4 (white dots). Traits above the dashed diagonal line show stronger parallel than non-parallel divergence across ecotype pairs. **(b)** Illustration of all seven linear traits measured: HDE – head depth at eye, HDO – head depth at operculum, HL – head length, PFL – pectoral fin length, ED – eye diameter, ML – maxilla length, LJL – lower jaw length. **(c)** Mean trait-values are plotted for each benthivorous-planktivorous and piscivorous-planktivorous ecotype pair, with means for sympatric pair being connected by a line. These reaction norms are colour coded blue and red highlighting the decrease or increase of trait values, respectively, between benthivorous (bn) or piscivorous (pisc) and planktivorous (pl). **(d)** Principal components plot based on all seven linear traits showing the centroid for each ecotype (N=1,329 individuals), with centroids of sympatric ecotypes connected by trajectories. Points are coloured by ecotype: bn – benthivorous, pl – planktivorous, pisc – piscivorous, pisc-s – small-piscivorous, insct – insectivorous.

For a more comprehensive analysis of parallelism that synthesizes across traits, we used a phenotypic trajectory approach to quantify the direction and magnitude (length and direction of trajectories) of phenotypic divergences across replicate sympatric ecotypes (Adams & Collyer 2009; Stuart et al. 2017). The length of these trajectories (L) describes the magnitude of divergence, and the angle ! between trajectories their direction through multi-trait space (see (Stuart et al. 2017; Bolnick et al. 2018) for details). Thus, the difference in trajectory length (ΔL_p_) and the direction of trajectories (*θ*_p_) defines the extent of multivariate phenotypic parallelism; completely parallel ecotype-pairs are those diverged to the same extent (ΔL_p_ not different from zero) and in the same direction (*θ*_p_ angle not different from zero) (Stuart et al. 2017; Bolnick et al. 2018). Using this approach, we found that the direction of phenotypic change between ecotype pairs was highly variable across replicates (Fig. 2d, Supplementary Fig. S1). Angles ranged from highly parallel (e.g. *θ*_p_ = 14.2°, P = 0.27 between the benthivorous-planktivorous ecotypes from Kamkanda and Davatchan) to strongly antiparallel (e.g. *θ*_p_ = 133.5°, P = 0.001 between the benthic-pelagic ecotypes from Awe and na Sealga) (Supplementary Fig. S1c, Supplementary table S3). The average angle between phenotypic trajectories across replicated ecotype pairs was *θ*_p_ = 58.95° ± 31.54 s.d. (standard deviations), meaning that trajectory directions differed, on average, widely across ecotype pairs, although they were smaller (more parallel) than reported e.g. for lake-stream sticklebacks (Stuart et al. 2017) (Supplementary table S3). Several ecotype pairs were significantly parallel in their direction of phenotypic divergence (Supplementary table S3). In general, angles between replicated ecotype pairs (e.g. benthivorous-planktivorous pair vs. benthivorous-planktivorous pair) were more similar than across nonreplicated ecotype pairs (e.g. benthivorous-planktivorous pair vs. piscivorous-planktivorous pair), which were on average orthogonal to each other (*θ*_p_ = 93.48° ± 18.94 s.d.; Wilcoxon rank sum test: P<0.001).

Compared to the direction of change, ecotype pairs were more similar in the magnitude of divergence (mean ΔL_p_ = 0.033 ± 0.021 s.d.), suggesting the extent of divergence between sympatric ecotypes is potentially more predictable and less influenced by contingent and stochastic (environmental or evolutionary) factors.

Some ecotype pairs were significantly parallel in both direction and magnitude, such as the piscivorous-planktivorous ecotypes from Kiryalta-3 and Kalarskii-Davatchan (*θ*_p_ = 26.8°, P = 0.35; ΔL_p_ = 0.0198, P = 0.311) and the benthivorous-planktivorous ecotypes from Kamkanda and Davatchan (*θ*_p_ = 14.2°, P = 0.27; ΔL_p_ = 0.02, P = 0.049). In general, Siberian ecotype-pairs, particularly the piscivorous-planktivorous ecotypes, showed a higher degree of parallelism (mean *θ*_p_ = 46.6° ± 21.3 s.d./ mean ΔL_p_ = 0.035 ± 0.020 s.d.) than ecotype-pairs from the Atlantic lineage (*θ*_p_ = 69.4° ± 35.6 s.d./ mean ΔL_p_ = 0.036 ± 0.021 s.d.; Supplementary Table S3). However, significant parallelism was not only restricted to within lineage comparisons, as the benthivorous-planktivorous ecotype pairs from Dughaill and Kamkanda were highly parallel in both, direction and magnitude of divergence (*θ*_p_=24.4°, P=0.07; ΔL_p_=0.0152, P=0.244).

Overall, these results indicate that ecologically replicated Arctic charr ecotypes show significant phenotypic parallelism in direction and magnitude. This is particularly strong for some individual traits relevant for foraging and swimming (Klemetsen et al. 2002), such as eye diameter and fin length, as highlighted by significant effect sizes and parallel trait trajectories. Parallelism in traits associated with ecological niche are considered a hallmark of repeated environmental adaptation (Schluter 2001) but robust empirical examples have been rare (Elmer et al. 2014; Kusche et al. 2015; Oke et al. 2017). In contrast to particular traits, we find substantial (non)-parallelism exists in the direction and magnitude of overall phenotypic divergence among replicated ecotype-pairs, but that this is not hierarchically structured across populations and lineages. These deviations from significant eco-morphological parallelism could be due to differences in genetic, demographic and environmental conditions (Stuart et al. 2017; Kusche et al. 2015; Oke et al. 2017; Collyer & Adams 2013).

### Evolution via non-parallel genomic routes

Based on the observed variation in extent of eco-morphological parallelism across replicated ecotype pairs we expected variation in the extent of population genetic parallelism. However due to the significant prevalence of eco-morphological parallelism some genetic parallelism is expected, which we predicted to be higher within lineages, particularly within the Siberian lineage, than across. To test this, we examined the repeatability of genome-wide patterns of differentiation and of putatively adaptive loci.

Principal components analysis based on a global SNP dataset of all individuals (N=12,215 SNPs; N=630 individuals) revealed that sympatric ecotypes strongly clustered by lake and river catchment (Gordeeva et al. 2015; Wilson et al. 2004), differed widely in their degree of genetic differentiation, and mostly split along independent principal components (Fig. 3a; Supplementary Fig. S2). This pattern suggests that ecotype pairs evolved independently from each other by different population genomic processes following the colonization of newly formed postglacial lakes (McVean 2009). Indeed, ecotype pairs did not share more outlier loci (top 5%-Fst outlier loci; Supplementary Fig. S3a-c, Supplementary table S4), or contigs containing outlier loci, than expected by chance in pairwise comparisons (Fig. 3b,c), indicating that genomic responses to selection differ across ecotype-pairs within and across lineages. Trajectory angles of neutral (N_Total_=7,179) and outlier (N_Total_ =986) loci did not differ between replicated ecotype pairs nor across divergent ecotype pairs, supporting the view that neutral patterns of divergence and responses to selection are mostly lake specific (Supplementary Fig. S3d). Similarly, loci putatively under selection within each lineage, (PCadapt: N_sel,ATL_=599; N_sel,SIB_ =666 SNPs; Supplementary Fig. S3) were not shared more often than expected by chance between the lineages (Supplementary Fig. S3e,f; N_shared_=18, P=0.51). Overall, these results show that replicated ecotype pairs have evolved repeatedly and independently using different genomic routes.

**Figure 3.**
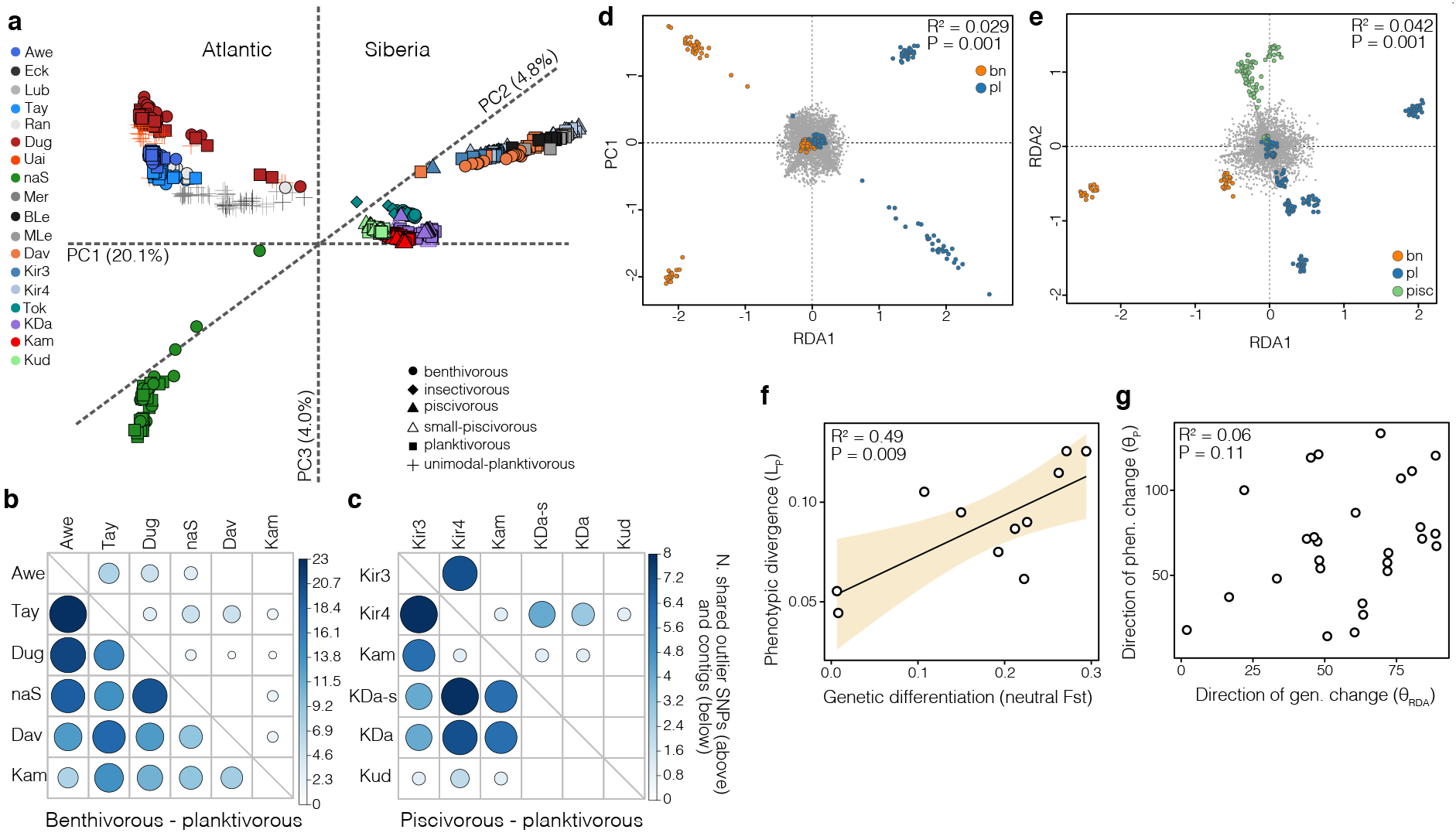
(Non)-parallelism in differentiation and ecotype-association. **(a)** 3-dimensional principal components plot for all individuals (N=630) based on 12,215 SNPs. Lake abbreviations are explained in Supplementary table S1. **(b,c)** Sharing of outlier SNPs and contigs containing outlier SNPs across replicated benthivorous-planktivorous **(b)** and piscivorous-planktivorous **(c)** ecotype pairs. The colour and size of the dots illustrates the number of shared SNPs or contigs. None of the pairwise comparisons are significant based on permutation results (see methods). **(d,e)** Results of the constrained redundancy analysis (RDA) for the Atlantic and Siberian lineage, showing varying levels of separation between ecotypes. The RDA significantly separates ecotypes after correcting for the effect of lake (results of ANOVA shown in plot). **(f)** Significant correlation between the degree of neutral genetic differentiation (neutral Fst) and phenotypic divergence (L_p_), with the shaded area highlighting the 95^th^ confidence interval (CI) (N=11). **(g)** The direction of allele frequency change of ecotype-associated loci (θ_RDA_) did not explain differences in the direction of phenotypic change (θ_p_) across ecotype pairs (N= 27).

As ecotypes show some parallelism in overall phenotypic divergence and particularly in distinct eco-morphological traits, we also expected significant allele frequency parallelism in ecotype-associated loci within and across lineages. We identified 217 and 303 loci significantly associated (z-score > 2) with ecotype divergence based on redundancy analyses within the Atlantic and Siberian lineages, respectively (Fig. 3d,e), which explained 2.9% and 4.2% of the overall variation between ecotypes within each lineage. This suggests higher levels of parallelism in the Siberian lineage. Of these, eight were shared across lineages, which is more than expected by chance (*χ*^2^-square; P<0.001; Supplementary table S5). In contrast, allele frequency patterns of ecotype-associated loci were mostly non-parallel across ecotype pairs, when within lineages, based on trajectory analysis of principal component scores (Supplementary Fig. S3f). The direction and magnitude of allele frequency change differed widely across populations, ranging from highly parallel (piscivorous-planktivorous from Kiryalta-3 and Kiryalta-4) to non-parallel (piscivorous-planktivorous from Kamkanda and Kiryalta-4). As expected, levels of parallelism in allele frequencies were more similar between replicates of the same type of ecotype pair (Mean θ_RDA,bn-pl_= 46.9° ± 16.0 s.d./ mean ΔL_RDA,bn-pl_=26.5 ± 19.2 s.d.; mean θ_RDA,pisc-pl_ 49.1° ± 18.7 s.d./ mean ΔL_RDA,pisc-pl_= 20.0 ± 12.6 s.d) than across different ecotype pairs (Mean θ_RDA,across_ 82.6° ± 16.6 s.d./ mean ΔL_RDA,across_= 30.2 ± 20.8 s.d.), supporting the role of these loci in ecotype divergence. Overall, these findings demonstrate that at least part of the genetic variation associated with ecotype is shared across ecotype pairs and also across lineages, including a potential core set of genes involved in ecotype divergence in the Atlantic and Siberian lineage.

The variability in the genetic basis of the ecotype and extent of genetic differentiation (neutral and outlier loci) across lakes and lineages might impact the extent of phenotypic parallelism (Stuart et al. 2017). Thus, we tested for correlations between the directions and magnitudes of genetic and phenotypic trajectories. We hypothesized that stronger neutral genetic differentiation, indicating less gene flow between sympatric ecotypes, would be associated with stronger phenotypic divergence, as predicted by theory (Shafer & Wolf 2013) and suggested by studies in other systems (Stuart et al. 2017; Nosil et al. 2009; Elmer et al. 2010; Manousaki et al. 2013). Neutral Fst between ecotypes was strongly correlated with the magnitude of phenotypic divergence (Fig. 3f; neutral F_st_ ~ L_p_: R^2^ = 0.493, P = 0.009), indicating that the degree of genetic differentiation potentially predicts the extent of phenotypic divergence. However, differences in the directions of the genetic trajectories (θ_Gn, Go_ and *θ*_RDA_), which show variation in the direction of allele frequency changes across ecotype pairs, were not correlated with differences in the directions of phenotypic trajectories (*θ*_p_) (Fig. 3g). This demonstrates that variation in the direction of genetic differentiation is not underlying variation in phenotypic parallelism in Arctic charr (for example, R^2^ = 0.06 in Fig. 3g). This is consistent with our result that there is little sharing of outlier or ecotype-associated SNPs across ecotype pairs (Fig. 3b,c).

Thus, although the extent of background genetic differentiation is correlated with the magnitude of phenotypic divergence, potentially through differences in the extent of gene flow, the observed variation in the direction of phenotypic divergence is most likely not explained by patterns of genetic variation in neutral nor putatively adaptive loci. In fact, there is relatively low sharing of outlier loci across population replicates, even between nearby lakes. This contrasts with findings from other postglacial fishes, for example sticklebacks (Stuart et al. 2017), where repeated divergences from a shared ancestral population facilitates rather high levels of outlier sharing, but agrees with findings in stick insects^17^, where an enrichment of unique divergences was identified between ecotype pairs from a shared ancestral population.

### Heterogeneous patterns of genetic co-ancestry

The observed heterogeneity in patterns of genetic differentiation suggests complex and varying evolutionary histories underlying the evolution of sympatric ecotype pairs. Thus, we investigated patterns of shared ancestry and secondary gene flow within and across lake populations and compared them to patterns of phenotypic and genetic differentiation. We predicted that populations that were more similar in their genomic background would show more similar patterns of genetic divergence of their ecotype pairs (Stuart et al. 2017).

Genetic co-ancestry analyses showed that ecotypes mostly formed genetically distinct clusters, with sympatric ecotypes having higher levels of shared genetic background, but also revealed clustering of allopatric ecotypes from lakes in the same catchment (Fig. 3a, Fig. 4a,b; Supplementary table S1, Supplementary Fig. S2). These patterns were also supported by a phylogenetic split network (Supplementary Fig. S4) and admixture analysis (Fig. 4c,d), suggesting different levels of genetic sharing and isolation between sympatric ecotypes and across lakes. In only two cases, sympatric ecotype pairs formed polyphyletic genetic clusters, with one ecotype being genetically more similar to the ecotype from a neighbouring lake than its sympatric pair (piscivorous ecotypes in Kiryalta-3 and Kiryalta-4, planktivores from Dughaill and Uaine; Fig. 5a; Supplementary Fig S4). Sympatric ecotypes pairs also differed widely in their degree of genetic differentiation (mean Fst ranging from 0.011 to 0.329), with differentiation between sympatric ecotypes in some cases being higher than between allopatric populations (Supplementary Fig. S5). This consistent history of divergences within lakes was further supported by near complete mitochondrial haplotype sharing across sympatric ecotypes in most populations and in some cases across nearby lakes (Supplementary Fig. S6).

**Figure 4.**
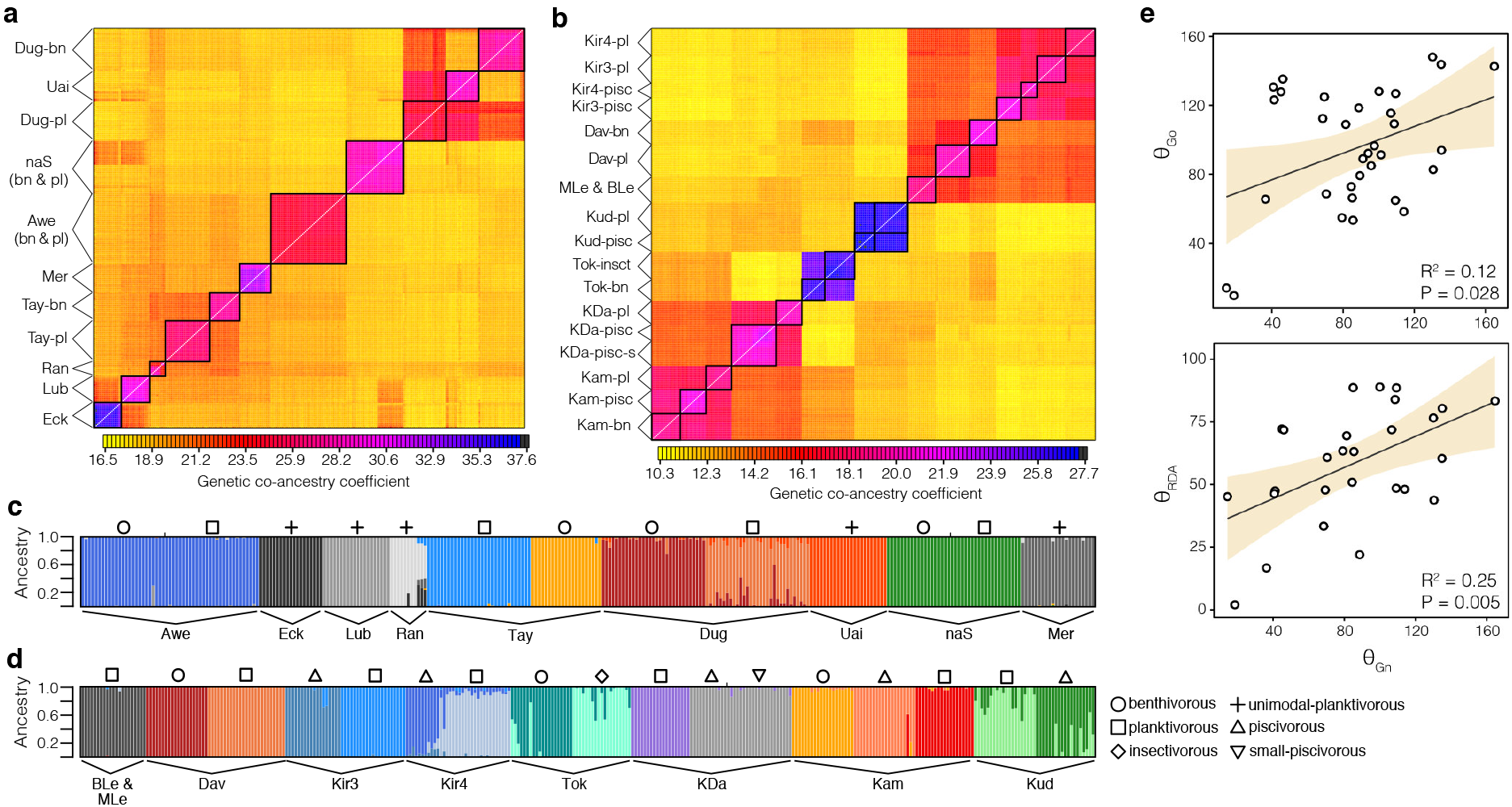
Population genetic structure. **(a,b)** fineRADstructure results showing co-ancestry coefficients between individuals from the **(a)** Atlantic and **(b)** Siberian lineage. Ecotypes or lake populations that form discrete genetic clusters are enclosed in black boxes. Note the high genetic co-ancestry across lakes within the same catchment, such as **(a)** Dughaill (Dug) and Uaine (Uai) or (b) Kiryalta-3 (Kir3), Kiryalta-4 (Kir4) and Davatachan (Dav). **(c,d)** *Admixture* plots showing the genetic ancestry for all individuals from the **(c)** Atlantic and **(d)** Siberian lineage for K=11 and K=16, respectively. Ecotypes are marked by symbols above each cluster. **(e)** Correlations between similarities in neutral allele frequency patterns (θ_Gn_) and allele-frequency patterns of outlier SNPs (θ_Go_, N=34) and ecotype-associated SNPs (θ_RDA_, N=27) across Arctic charr ecotype pairs. The shaded area shows the 95^th^ CIs around the regression lines.

In general, contemporary neutral patterns of genetic divergence, represented by angles between neutral genetic trajectories, correlate with differences in the direction of outlier SNPs (Fig. 4e; θ_Gn_ ~ θ_Go_: R^2^=0.119, P = 0.028) and ecotype-associated SNP trajectories (Fig. 4e; θ_Gn_ ~ θ_RDA_: R^2^=0.245, P=0.005). Thus, populations that are genetically more similar at neutral SNPs, i.e. having higher shared ancestry, are also more similar in allele frequency divergence in differentiated and putatively adaptive loci.

Similarities in genetic background are expected to lead to more similar genetic responses to selection, e.g. through higher proportions of shared genetic variation (Conte et al. 2012). In general, our findings show low levels of sharing of putatively adaptive genetic material across lakes. This result is in contrast to other well studied examples of postglacial divergences in fishes, such as normal-limnetic lake whitefish (Rougeux et al. 2017) and benthic-limnetic stickleback fishes (Jones et al. 2012), in which multiple invasions of ecotype-specific adaptive material facilitated divergence into sympatric pairs. However, we find that populations with more similar neutral background divergence are also more similar in putatively adaptive allele frequency trajectories. Together, this is consistent with repeated and independent divergences between ecotypes within lakes from standing genetic variation rather than introgression of adaptive genetic material across populations.

### Variable and non-parallel evolutionary histories underlie parallel phenotypic divergence

Contemporary patterns of shared ancestry and genetic differentiation indicate a role for complex historical gene flow and diversification within and across populations. To test for historical gene flow and introgression events across lakes that potentially led to non-independent patterns of eco-morphological divergence, we inferred putative 22’secondary gene flow events using *f4-* and *f3*-statistics implemented in *Treemix.* Overall, these analyses indicate complex histories of gene flow across catchments and lakes in both lineages (Fig. 5a, Supplementary Fig. S4a-c, Supplementary table S6) that might have contributed to the complex patterns of shared ancestry. However, our results suggest that these historical migration events do not explain the repeated divergences of ecotypes within lakes e.g. through the introgression of adaptive genetic material (Richards et al. 2018), as migration events appear random across ecotypes and lakes. Though, more detailed future investigations are needed to fully exclude the possible role of adaptive introgression.

**Figure 5.**
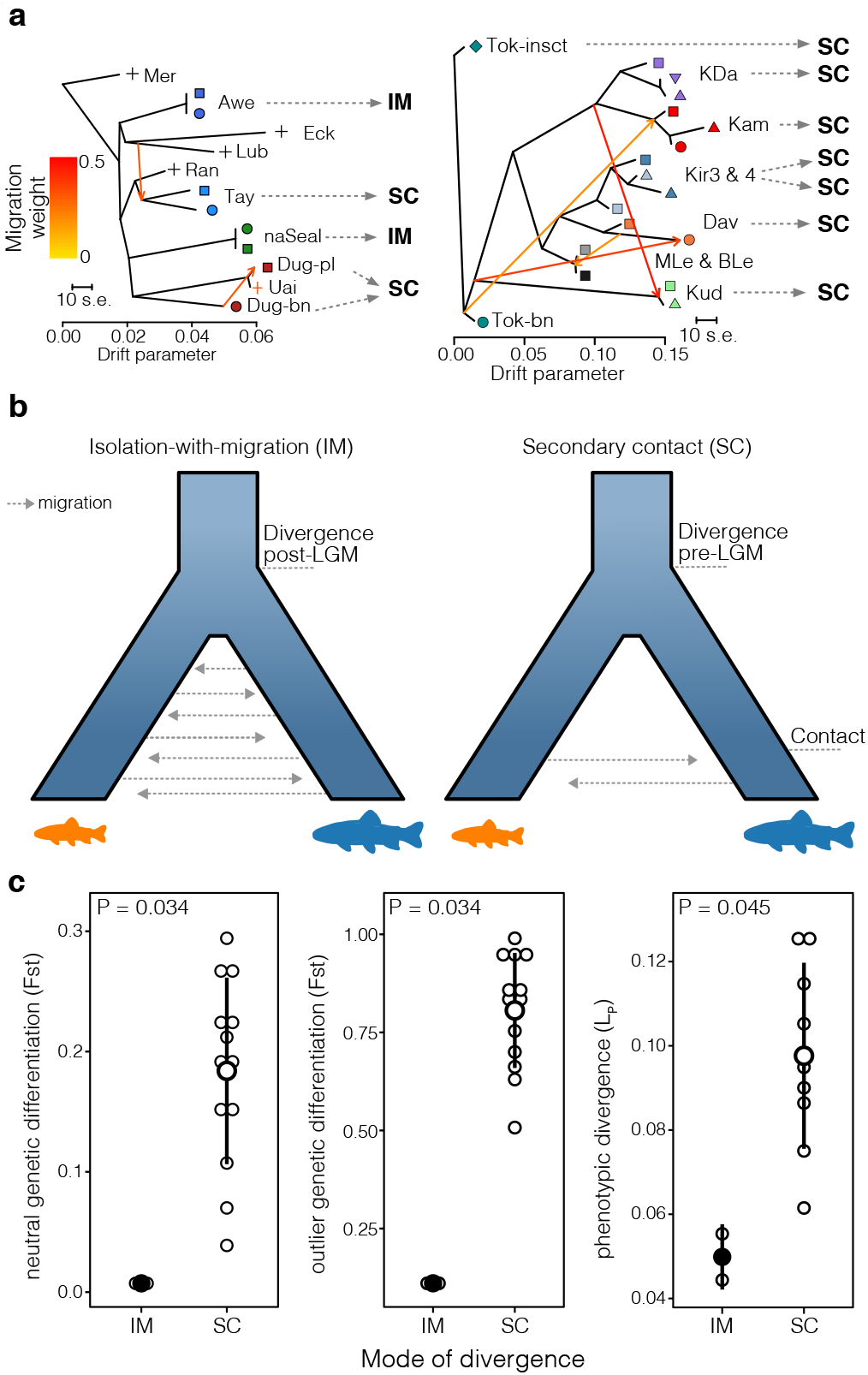
Secondary gene flow and evolutionary histories. **(a)** Allele-frequency based maximum-likelihood trees showing the most likely phylogenetic relationships across lakes and ecotype pairs by lineage, including two and four fitted migration events in the Atlantic and Siberian lineage, respectively. Migration events are shown as arrows coloured by migration weight. The most likely evolutionary history for each ecotype pair is shown behind the dotted arrows, and the models of evolutionary histories are illustrated in panel **(b)** and Supplementary Fig. S8. **(b)** Illustrations of the Isolation-with-migration (IM) and secondary contact (SC) models. **(c)** Differences in the magnitude of phenotypic divergence, and neutral and outlier genetic differentiation between ecotype pairs evolved under different evolutionary histories (IM vs SC) (N_IM_=2; N_sc_=9-13).

To understand the effect of historical demography on contemporary genetic patterns and the extent of parallelism, we modelled the evolutionary history of all sympatric ecotypes. We used coalescence simulations implemented in *fastsimcoal2* (Excoffier et al. 2013) for two-population and three-population models (Supplementary Fig. S7). Most ecotype pairs (11 out of 13 pairs) likely evolved via historical secondary contact that resulted in the admixture of those ancestral populations (Fig. 5b; Supplementary Fig. S8, Supplementary table S7). Considerable variation existed in initial divergence times between ancestral populations (910 – 16,690 generations ago), timing of secondary contact (56 – 3,783 generations ago), and the arising of contemporary sympatric ecotypes (Supplementary Fig. S8). The extent of admixture between ancestral populations upon secondary contact also varied widely across lakes, with genome-wide proportions from 0.003 – 1.0 (Supplementary Fig. S8). These model results also suggested that ancestral populations in Siberia were strongly isolated during the last glacial maximum (LGM; 10-20 kya), while weak gene flow in the Atlantic populations was detected during the LGM (Supplementary Fig. S8). In combination with these complex and divergent evolutionary histories, genetic and ecological data suggest that most ecotypes likely then diverged in sympatry (Gordeeva et al. 2015), facilitated by genetic variation that arose in allopatry across ancestral populations (Garduño-Paz et al. 2012; Rougeux et al. 2017). In two ecotype pairs, we found evidence for divergence-with-gene-flow of ecotypes arising in sympatry after the LGM (isolation-with-migration model; T_DIV,Awe_=1245 generations/3735 years, T_DIV,nas_ =4795 generations/14385 years) (Fig. 5b, Supplementary Fig. S8), consistent with incipient sympatric speciation (Coyne & Orr 2004).

We further tested whether these differences in historical demographic parameters are associated with differences in the extent of genetic differentiation and thereby potentially the magnitude of phenotypic divergence in ecotype pairs. Indeed, we found that ecotype pairs that most likely diverged under secondary contact showed a higher degree of neutral genetic differentiation (ΔFst_neut_=0.186, P=0.034) and differentiation in outlier loci (ΔFst_out_=0.731, P=0.034) compared to ecotype pairs that diverged under gene flow (Fig. 5c). This was also true for the extent of overall phenotypic divergence (Fig. 5c; ΔL_p_=0.047, P=0.045). The extent of neutral genetic and phenotypic divergence was not significantly correlated with divergence time, timing of secondary contact, or the magnitude of contemporary migration rate (Supplementary Table S8), although neutral Fst and neutral divergence (L_Gn_) tended to increase with divergence time and decrease with increasing migration rate (L_Gn_ ~ T_DIV_: R^2^=0.199, P=0.071; neutral Fst ~ Nm: R^2^=0.204, P = 0.052). However, differences in divergence time did explain differences in the degree of genetic differentiation in outlier loci (L_Go_ ~ T_DIV_: R2=0.334, P=0.0226), suggesting that genetic differentiation in putatively adaptive loci builds up over time.

These results indicate that the mode of divergence (secondary contact or isolation-with-migration) had a stronger impact on contemporary patterns of parallelism between ecotype pairs than did demographic parameters (i.e. time since divergence). Specifically, ecotype divergence following secondary contact with admixture was associated with higher phenotypic and genetic differences. Thus, evolutionary and demographic history has a measurable and predictable effect on parallelism in the magnitude of divergence.

### Parallel divergence in gene expression

Given the relatively low parallelism in genomic patterns compared to parallelism in phenotype, we hypothesized that regulatory variation would show parallelism among ecotypes due to integrated effects of plasticity and genetically-mediated expression. Therefore, we analysed genome-wide gene expression between ecotypes from a subset of five lakes (N=44 individuals, 44,102 genes; Supplementary table S1), and compared differential expression, co-expression modules, and biological pathways.

Similar to population genetic patterns, we found a continuum of divergence in gene expression across ecotype pairs (Fig. 6a; Supplementary Fig. S9a). Contrary to the genomic analysis, though, gene expression patterns were highly similar across lakes and ecotype pairs, with ecotype explaining most of the expression variation along PC1 (Fig. 6a; η^2^_Eco,PC1_ = 0.80, P Δ 0.001) and more than the non-parallel interaction terms (non-significant except for PC4) for PC2 to PC4 (Supplementary Fig. S9b). Differentially expressed genes (DEGs) were shared between replicated ecotype pairs significantly more often than expected by chance (Fig. 6b), indicating highly parallel divergences in gene expression. We identified four genes (Fig. 6c; *ABCC8, NDPK, ALDOA, uncharacterized protein LOC100194706)* that were consistently differentially expressed in five out of seven ecotype pairs (Fig. 6c). These genes are known to be associated with growth rate, body size, and metabolism (Morbey et al. 2010; Palstra et al. 2014; Flanagan et al. 2017), making them strong candidates for underlying ecotype divergence in Arctic charr. Additionally, 19 differentially expressed genes were shared across four ecotype pairs, which included several haemoglobin subunits (Supplementary Fig. S9c).

**Figure 6.**
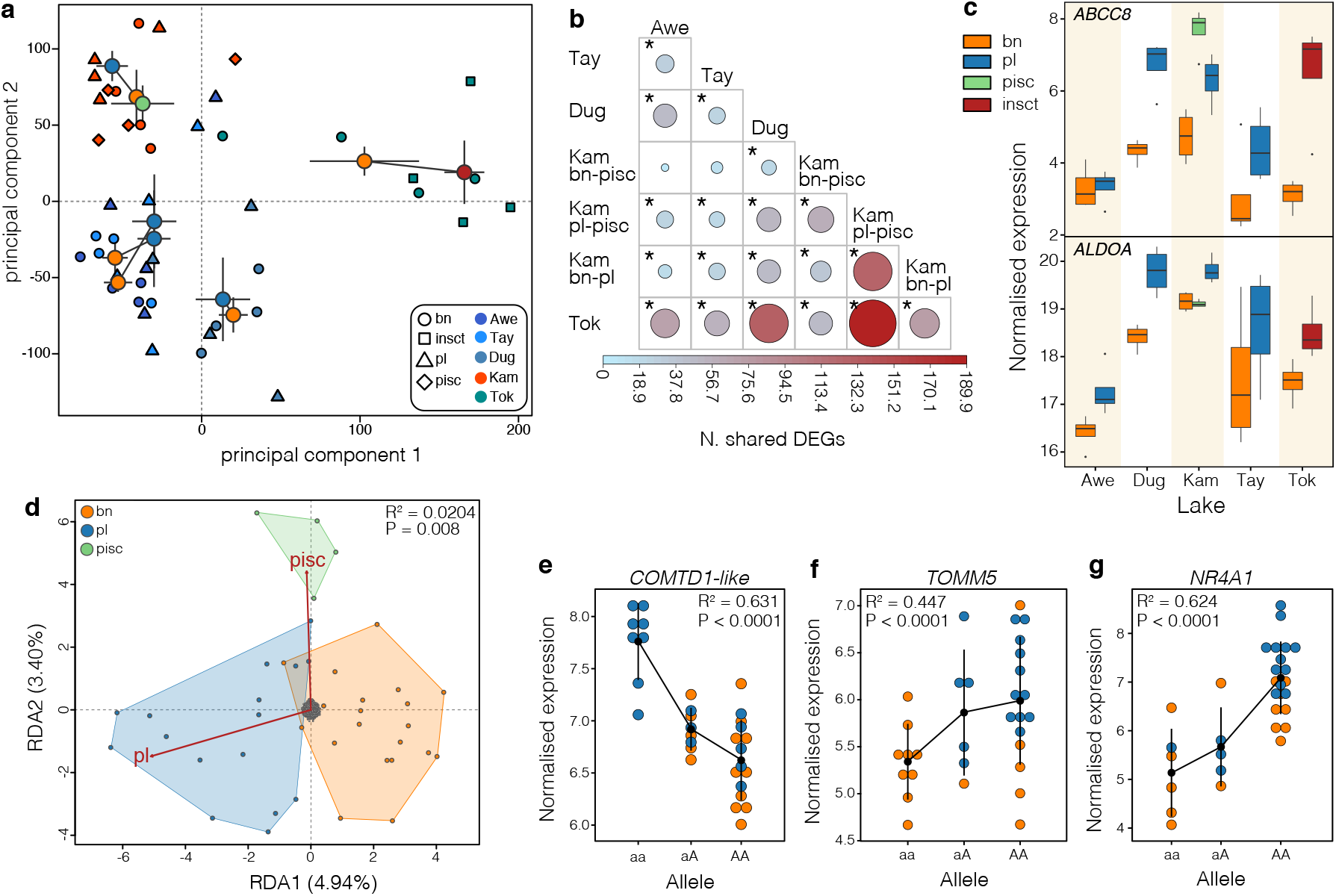
Parallelism and divergence in gene expression. **(a)** Principal components plot based on r-log transformed gene expression data (N=30,849 transcripts). Individuals are shown by individual points shaped by ecotype and coloured by lake of origin. Centroids for each ecotype are shown including standard error and coloured by ecotype (blue – planktivorous, orange – benthivorous, green – piscivorous, red – insectivorous). Centroids of sympatric ecotypes are connected by a line. **(b)** Sharing of differentially expressed genes with the extent of sharing weighted by colour and circle size. Significant comparisons are highlighted with an asterisk. **(c)** Expression of two genes *(ABCC8* and *ALDOA)* significantly differentially expressed in 5 out of 7 ecotype pair comparisons. **(d)** Biplot of the gene expression based constrained redundancy analysis showing significant separation of ecotypes after correction for lake and lineage, with benthivorous (orange area) and planktivorous (blue area) separating along RDA1 (4.94%) and the piscivorous (green area) splitting off along RDA2 (3.40%). (eg) Examples of *cis*-eQTL in **(e)** COMTDl-like, **(f)** *TOMM5* and **(g)** NR4A1, showing how the expression of these ecotype-associated genes differs with genotype across individuals (points) and ecotypes. The black dot and range show the mean expression per allele and standard deviation.

Using a redundancy analysis, we identified 2,921 genes that have expression correlated with ecotype (z-score > 2), which explain 2.04% of the variation in gene expression (P = 0.008), after correcting for the effect of lake and lineage (Fig. 6d). This included eight out of the 23 DEGs that were significantly shared across at least four ecotype pairs, supporting their importance for ecotype divergence. Of these ecotype-associated differentially expressed genes, 25 (0.85%) were associated with cis-regulatory variation (cis-eQTL; FDR < 0.1), indicating that their expression divergence between ecotypes is at least partially genetically determined (Supplementary table S9). The most significant ecotype-associated *cis*-eQTLs included the COMTD1-like gene, an enzyme associated with muscle mass in humans (Ronkainen et al. 2008), *TOMM5*, a crucial protein involved in mitochondrial protein import, and *NR4A1* which is involved in gene expression regulation (Fig. 6e-g; Supplementary table S9). Due to the relatively low sample size we likely only detected the strongest *cis*-eQTLs and underestimated the true number of genes with genetically-mediated differential expression. Furthermore, we identified 13 modules of co-expressed genes (N = 806 genes) that were correlated with ecotype divergence for benthivorous-planktivorous ecotype pairs across lakes and lineages (Supplementary figure S9e, Supplementary table S10). Of these, 162 genes were also ecotype-associated in the redundancy analysis, further indicating the importance of expression changes in gene networks and suggesting parallel changes in regulatory networks across lakes and lineages.

Overall, ecotype-associated genes (from GEx-RDA) were involved in a range of biological processes, including cell cycle regulation (GO:0007049, Fold-enriched = 32; FDR < 0.001), chromosome organization (GO:0051276, Fold-enriched = 11; FDR < 0.001), chromatin organization (GO:0006325, Fold-enriched = 19; FDR = 0.059) or microtubule-based processes (GO:0007017, Fold-enriched = 19; FDR = 0.013) (Supplementary table S11), which are processes functionally associated with growth, cell differentiation and gene regulation.

In contrast to genomic changes, differences in the direction and magnitude of gene expression divergence between sympatric ecotypes explained the differences in patterns of phenotypic divergence across Arctic charr ecotype pairs (Fig. 7). Differences in the magnitude of gene expression divergence (ΔL_GEx_) were significantly positively correlated with differences in the magnitude of phenotypic divergence (–L_p_) in comparisons across all ecotype pairs (–L_p_ ~ ΔL_GEx_: R^2^= 0.367, P=0.0371; Fig. 7a). Sympatric ecotype pairs that were more similar in the direction of gene expression change in ecotype-associated genes (θ_canGEx_) tended to be also more similar in the direction of phenotypic change (*θ*_p_) (*θ*_p_~θ_canGEx_: R^2^=0.236, P=0.0878; Fig. 7b). This suggests that divergence in gene expression plays an important role in these adaptive diversifications, and that variation in gene expression putatively explains the variation in phenotypic parallelism seen in Arctic charr; this effect holds not only across lake replicates but also across evolutionarily distinct lineages.

**Figure 7.**
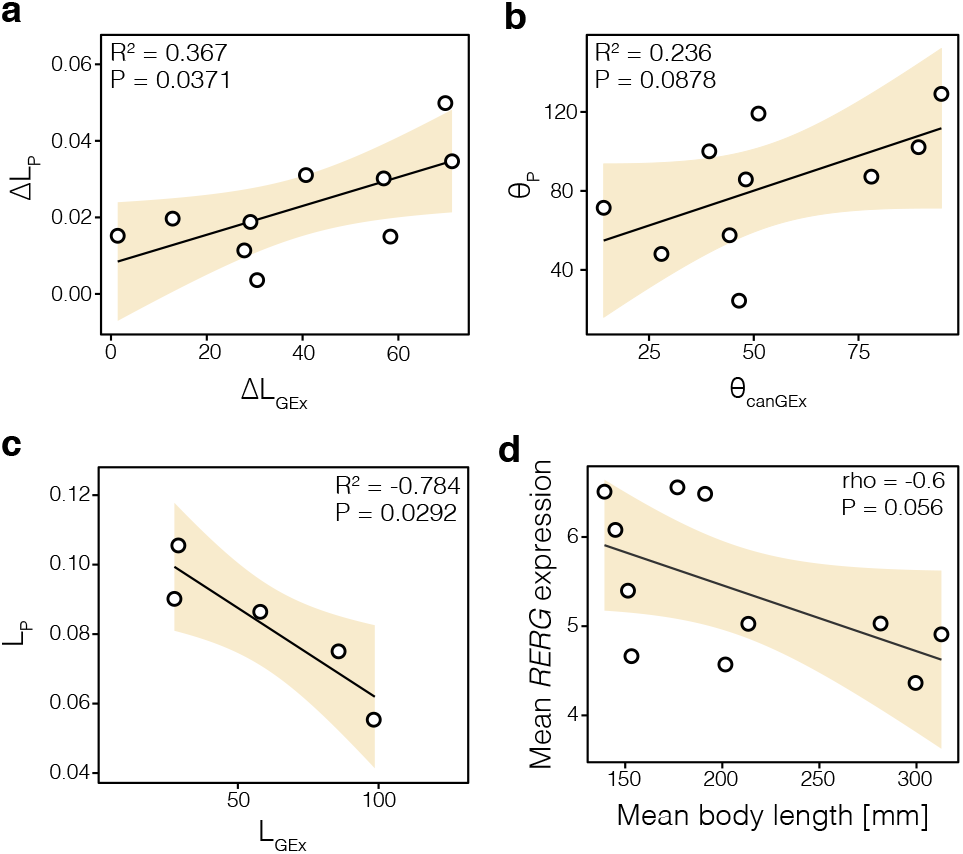
Correlation of gene expression and patterns of phenotypic divergence. **(a)** Differences in the magnitude of phenotypic divergence (ALP) are significantly correlated with differences in the degree of phenotypic divergence (ALGEx) between sympatric Arctic charr ecotypes (N=10). **(b)** The direction of expression trajectories of ecotype-associated genes (θcanGEx) tends to be correlated with the direction of the phenotypic trajectories (θP) (N=10). **(c)** Negative correlation between the absolute magnitude of expression divergence (LGEx) and phenotypic divergence (LP) (N=5). (d) The mean expression of *RERG* is negatively correlated with the mean body length of ecotypes across lakes and lineages (N=11).

One might expect that ecotypes that are more strongly diverged in gene expression might also be phenotypically more diverged. Contrary to this expectation, we observed that gene expression divergence between sympatric ecotypes is negatively correlated with phenotypic divergence (L_p_~L_GEx_: R^2^= 0.784, P=0.0292; Fig. 7c), with more phenotypically diverged ecotype pairs (which are also genetically more diverged) showing less gene expression divergence. We hypothesize that increased divergence in gene expression is partially a plastic reinforcement mechanism underlying adaptation despite lower genetic divergence. Additionally, we found that the expression of *RERG*, a growth inhibitor associated with ecotype-divergence, tended to be negatively correlated with the mean body size of ecotypes (Spearman’s rho = −0.6, P = 0.056), potentially explaining in part the huge diversity in body length across Arctic charr ecotypes (Fig. 7d).

In summary, we find that gene expression, which is demonstrably both heritable and plastic, is likely an important mechanism for the replicated and parallel phenotypic diversification of Arctic charr ecotypes, contributing to differences and similarities, even across divergent evolutionary lineages. We find sets of genes that are biologically relevant based on human and model organism studies (Morbey et al. 2010; Palstra et al. 2014; Flanagan et al. 2017; Ronkainen et al. 2008) and are associated with traits that differ between these and other fish ecotype divergences, such as haemoglobins (Supplementary figure S9c) (Evans et al. 2012; Filteau et al. 2013). This is consistent with the transcriptome being a flexible and rapid response to selection (Schneider & Meyer 2017) that can facilitate convergent evolution across ecotypes and species (e.g. (Gallant et al. 2014)).

### Predictability of genotypes and phenotypes in environmental context

The advantage of studying replicated natural populations is that one is able to infer the influence of the environment on patterns of diversification. To understand how environmental differences across lakes predict patterns of diversity and divergence, we examined correlations between ecosystem size (see methods), a proxy for ecological niche diversity within a lake, with phenotypic, genetic and gene expression patterns. Based on theory and previous studies (Recknagel et al. 2017; Kautt et al. 2018; Kusche et al. 2014), we expected that larger ecosystems allow for more diverse populations and stronger divergence, particularly in plastic traits.

In contrast to these expectations, we found that the extent of mean phenotypic divergence in individual phenotypic traits (mean Pst across traits) scaled negatively with ecosystem size (mean P_st_~Ecosystem size: R^2^=0.39, P=0.009; Fig. 8a), meaning that ecotype pairs in larger lakes are generally less diverged. On the other hand, the variance in phenotypic traits across ecotypes within a population increased with ecosystem size, suggesting that populations in larger and deeper lakes were more variable, although less diverged (mean trait variance~Ecosystem size: R^2^=0.73, P=0.001; Fig. 8b). However, phenotypic traits that showed low levels of parallelism (Fig. 2b) tended to be more strongly correlated with ecosystem size in their divergence and variance (Fig. 8c), indicating that these traits are potentially more plastic or adaptive to different environments, and thus more variable across different populations. As was found for phenotypic differentiation in non-parallel traits, variation in background genomic diversity across populations (measured as nucleotide diversity) was largely explained by differences in ecosystem size (Ecosystem size~n: R^2^=0.54, PΔ0.001; Fig. 8d). This suggests that larger lakes harbour more genetically diverse populations, which is in agreement with earlier research on phenotypic variability (Recknagel et al. 2017). However, patterns of genetic differentiation were in general not correlated to differences in ecosystem size (Ecosystem size~neutral Fst: R^2^ = 0.019, P = 0.28). Interestingly, we did not find any correlation between the extent of gene expression divergence and ecosystem size (Ecosystem size~L_GEx_: R^2^ = 0.13, P = 0.56), suggesting that habitat availability most likely does not predict gene expression divergence in Arctic charr. Divergence in gene expression is thus likely influenced by other factors, and not purely by ecosystem size. Thus, our results indicate that the extent of phenotypic divergence and variation, and therefore deviations from parallelism, can be largely explained by the interplay of environmental differences and differences in evolutionary history (mode of divergence; Fig. 5c). Similar predictable patterns have been found in Midas cichlids from Nicaraguan crater lakes (Kautt et al. 2018) and Canadian lake-stream sticklebacks (Stuart et al. 2017), demonstrating the impact of extrinsic conditions on the extent of parallel evolution.

**Figure 8.**
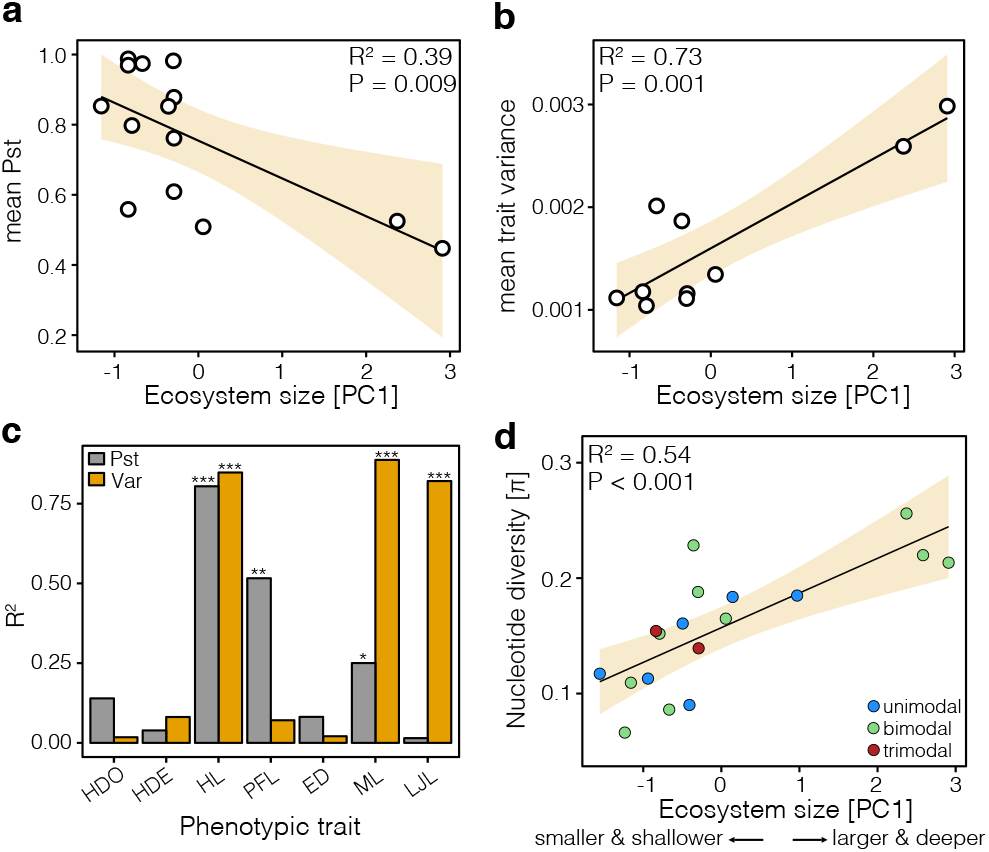
The correlation of ecological opportunity on divergence and diversity. **(a)** The mean degree of phenotypic differentiation (mean Pst) between sympatric ecotype pairs decreases with increasing ecosystem size in Arctic charr (N=14), whereas the **(b)** mean trait variance increases with ecosystem size (N=10). **(c)** Correlation coefficients (R^2^-values) for correlations individual traits differentiation (Pst) and variance (Var) with ecosystem size. Significant correlations are highlighted with asterisk (* for P < 0.05, ** for P < 0.01, and *** for P < 0.001). **(d)** The within-population nucleotide diversity (ecotypes combined by lake) positively correlates with ecosystem size. Points are coloured by the number of sympatric ecotypes within each lake.

## Conclusion

Our findings demonstrate significant parallel phenotypic changes in several traits associated with foraging, morphology, and ecological divergence in Arctic charr and a high repeatability of genome-wide gene expression. This is true within evolutionary lineages and geographically proximate populations but also across divergent evolutionary lineages separated by 60,000 years of evolution and large geographic distances. Furthermore, we identified shared loci associated with ecotype within and across lineages, most likely due to shared ancestral variation with repeated independent divergences. Taken together, this suggests that similar selection pressures, particularly on individual traits, led to the repeated evolution of similar phenotypes, and at least partially via similar genetic and gene expression pathways.

However, we also found significant deviations from parallelism, not only in combined phenotypes and specific morphological traits of Arctic charr ecotypes, but also in genotype patterns typical of response to selection, such as outlier loci and genomic regions (i.e. SNPs and contigs). Despite the significant sharing of ecotype-associated loci within and across lineages, allele frequency trajectories of those loci deviate from parallelism, which is explained by differences in genomic background across populations. These unpredictable, non-parallel responses are explained by different intrinsic and extrinsic factors. Specifically, we find that neutral genetic patterns of divergence and differences in demographic history are associated with the extent of phenotypic and genetic divergence between ecotypes as well as the genomic responses to selection, suggesting that genomic background and evolutionary history determine the extent of parallelism. Additionally, ecological opportunity within lakes, in combination with neutral genetic patterns, is significantly correlated with the extent of phenotypic differentiation and parallelism of ecologically relevant traits. Overall, our results strengthen the hypothesis that the interplay of environmental and demographic factors strongly shapes the extent of phenotypic and genetic parallelism (Stuart et al. 2017; Manousaki et al. 2013).

Despite large variation in genetic responses to selection, we found that gene expression ties together environmental and genetic effects, and that it is significantly associated with the magnitude and direction of phenotypic change. Specifically, gene expression divergence between sympatric ecotypes showed higher levels of parallelism than neutral or putatively-adaptive genetic variation, resulting in predictable phenotypic divergence between Arctic charr ecotypes even across distinct evolutionary lineages. Ecotype-associated gene expression was associated with a range of biological processes such as metabolism, growth, cell differentiation and gene regulation, potentially explaining the observed differences between sympatric ecotypes in morphology and body size, as well as swimming and foraging behaviour (Klemetsen et al. 2002; Hooker et al. 2016; Alekseyev et al. 2002). Some of the ecotype-associated variation in gene expression is genetically mediated, although plasticity most likely plays an important role in the eco-morphological divergence of Arctic charr (Adams & Huntingford 2004; Alexander & Adams 2004). We propose that gene expression is a bridge that facilitates parallel evolution of ecotypes despite variation in environment, genomic background and evolutionary history.

The evolution of replicated ecotypes has long fascinated naturalists and evolutionary biologists, as it indicates the predictable action of natural selection (Losos 2017). Our study demonstrates the role of quantifiable stochasticities and contingencies on the extent and consistency of eco-morphological parallelism. Thus, the repeated divergences of Arctic charr provide an example of how replaying the tape of life can lead to repeated and predictable outcomes, contrary to Gould’s predictions (Blount et al. 2018), but also illuminates the variable routes and mechanisms leading to parallel adaptations.

## Materials and Methods

### Arctic charr sampling

Fish were sampled from nine Scottish lakes (Atlantic lineage) and nine Transbaikalian lakes (Siberian lineage) (Lecaudey et al. 2018), between 1995 and 2015 using standard gill nets (Fig.1a). The sampled lake populations (we refer to all individuals within a lake as a population) contained different numbers and combinations of ecotypes (we refer to trophic specialists as ecotypes). We classified individuals into four ecotypes based on their primary diet (see (Jonsson & Jonsson 2001; Alekseyev et al. 2002; Garduño-Paz et al. 2012; Hooker et al. 2016; Gordeeva et al. 2015) for details): 1) Planktivorous — which feeds mainly on plankton throughout the year, 2) Benthivorous — which consumes a substantial proportion of benthic invertebrates, particularly during autumn and winter, 3) Piscivorous — which feeds mainly on other fish, 4) Insectivorous – which feeds largely on terrestrial insects and 5) unimodal-planktivorous – which represent non-diverged mainly plankton-feeding populations used as outgroups. After collection, we photographed the left side of each fish (Atlantic samples), or individual fish were preserved in formaldehyde for subsequent phenotypic analysis (Siberian samples). White muscle tissue (from underneath the dorsal fin and above the lateral line) and/or fin clips were taken for subsequent genetic and transcriptomic analysis and stored in absolute ethanol or RNAlater at −20°C.

### Analysis of linear morphological traits

Eco-morphological analysis was performed based on seven linear traits, on 345 individuals (N=4 populations) from the Atlantic lineage and 984 individuals from the Siberian lineage (Supplementary Table 1). Seven linear measurements and fork length were taken from photographs using ImageJ v. 1.50i (Schneider et al. 2012) for Atlantic samples or directly from formaldehyde fixated fish for Siberian samples (Fig. 2c) (Alekseyev et al. 2002): FL — fork length, HDO — head depth at operculum, HDE — head depth at eye, HL — head length, ED — eye diameter, ML — maxilla length, LJL — lower jaw length, PFL — pectoral fin length. Linear traits were chosen based on previous studies on eco-morphological divergence in salmonid fishes and their potential functional importance (Alekseyev et al. 2002; Siwertsson et al. 2013; Klemetsen et al. 2002; Adams & Huntingford 2002). Linear traits were correlated with body length, and therefore scaled to mean fork length, using the allometric formula as described in Siwertsson et al. (2013): log^10^ Y_i_ = log^10^ M_i_ + *b* * (log^10^ L_m_ – log^10^ L_i_); where Y_i_ is the corrected trait-value, M_i_ is the measured trait value, *b* is the slope (regression coefficient) of the regression of the trait value against fork length (L_i_) within each lake and ecotype, and L_m_ is the mean fork length of all fish within a lineage. The slope was calculated using population and ecotype as covariates. Size-adjusted measurements were used for all subsequent analyses. Principal component analyses (PCA) were used to uncover the major axes of phenotypic variation in the Atlantic and Siberian lineages, using the *ppca* approach in *pcaMethods* (R-package) to account for missing data.

### Analysis of phenotypic parallelism based on linear traits

To determine the contribution of parallel and non-parallel aspects to the overall morphological divergence of ecotypes, within and across populations, we used the ANOVA (multivariate analysis of co-variance) approach outlined by Langerhans and DeWitt (Langerhans & DeWitt 2004). ANOVAs (trait/PC ~ ecotype + lake + lineage + ecotype x lake + ecotype x lineage) were performed for both lineages combined using individual principal component (PC) scores (PC1 to PC4) and individual linear traits to test for the extent of parallel (ecotype effect) and non-parallel (ecotype x lake interaction (E x L); ecotype x lineage (E x Lin) interaction) phenotypic divergence of sympatric ecotypes across lakes and lineages, and the effect of the unique evolutionary history of each ecotype pair (lake effect and lineage effect) on phenotypic variation across lakes and lineages. We used the *EtaSq* function in *BaylorEdPsych* (R-package) to estimate the effect size (Wilk’s partial η^2^) of each model term for linear traits and principal component scores. Traits and PCs for which the ecotype term has the largest effect size are highly parallel between ecotypes across lakes and lineages. Those traits and PCs for which the ecotype term explains more variation than the interaction terms (which indicate non-parallel patterns of divergence in magnitude and/or direction), but not more than the lineage and lake terms, are to some degree parallel but are strongly influenced by differences in the evolutionary history between lake populations or lineages

Furthermore, we performed a complementary phenotypic trajectory analysis (PTA) (Collyer & Adams 2013) to quantify the level of parallelism and deviations from parallelism based on all linear traits combined. The PTA was conducted using the *traįectory.ana!yńs* function in *geomorph* (R-package). The significance of differences in trajectory directions (θ_p_: differences in the direction of phenotypic change) and trajectory lengths (ΔL_p_: differences in the magnitude of phenotypic change) was assessed using 1,000 permutations. Ecotype pairs were considered parallel if the angle between trajectories and the differences in trajectory lengths were not significantly different from zero (see Stuart et al. 2017; Bolnick et al. 2018 for details).

### Chromosome assembly and annotation of Arctic charr draft genome

We created a draft chromosome-level assembly of the Arctic charr genome based on an Arctic charr linkage map(Nugent et al. 2016) and synteny with the Atlantic salmon reference genome(Lien et al. 2016) based on *Chromosomer* analysis (Tamazian et al. 2016). The assembly was created using *All Maps* (Tang et al. 2015). We created a draft annotation using *GeMoMa* 1.4.2 (Keilwagen et al. 2016) and the quality of gene predictions was evaluated using *BUSCO* v.1.22 (Simão et al. 2015). Arctic charr and zebrafish *(Danio rerió)* orthologues were identified using *Orthofinder* (Emms & Kelly 2015) (see Supplementary Notes for details).

### DNA extraction and ddRADseq

DNA was extracted from fin clips and muscle tissue using the NucleoSpin Tissue kit (Macherey-Nagel), following the manufacturers recommendations. DNA quality and quantity were assessed using agarose gel electrophoresis and the Qubit Fluorometer with the dsDNA BR Assay (Life Technologies). ddRADseq libraries were prepared using a modified version of the Recknagel et al. (2015) ddRADseq protocol for Illumina sequencing platforms. Paired-end 75-bp sequencing was performed on the Illumina NextSeq500 platform at Glasgow Polyomics (University of Glasgow) at 3-4M read coverage per individual.

### Amplification, sequencing and analysis of the mitochondrial ND1 gene

The mitochondrial ND1 gene was amplified for 107 individuals (between 2 and 11 individuals per population) using the primer-pair B1NDF/B1NDF and PCR conditions as described in Schenekar et al. (2014). The PCR product was cleaned, and Sanger sequenced in both directions at *DNA Sequencing and Services* (MRC I PPU). Contigs were assembled from forward and reverse reads using *Sequencher* v.5.4 *(http://www.genecodes.com*) after removing low quality reads and trimming read ends. Reads for all individuals were aligned using *Muscle* in *MEGA* v.7 (Kumar et al. 2016) and trimmed to a common length. A TCS haplotype network was built in *POPART* (Leigh & Bryant 2015).

### Processing of ddRADseq data

Raw sequence data quality was assessed using *FastQC v.0.11.3* (*http://www.bioinformatics.babraham.ac.uk/projects/fastqc*). The *process_radtags* pipeline in *Stacks* version 1.46 (Catchen et al. 2011) was used for demultiplexing raw sequence data based on unique barcodes, quality filtering and read-trimming to 70bp of all libraries. Processed reads were aligned to the Arctic charr draft genome with *bwa mem* v.0.7.15 using a seed length of 25bp and the —M option. Reads with mapping quality <20 were removed using *samtools* v.1.6. We used *Stacks* v.1.46 and the *ref_map.pl* pipeline for building RAD-loci and SNP calling. The *populations* module was used to export genotype calls in VCF format for further filtering in *vcftools* v.0.1.15. We created three different datasets; a global dataset for the Atlantic and Siberian lineages combined, and separate datasets for each lineage. SNPs were retained when the following criteria were fulfilled: (i) present in at least 66% of all individuals within a population and 2/3 of all populations, (ii) global minor allele frequency (MAF) ≥ 0.05 or ≥ 0.01 for the global dataset, (iii) heterozygosity ≤ 0.5 (iv) in Hardy-Weinberg equilibrium (P > 0.05) in at least 2/3 of all populations and (v) with a minimum coverage of 6x. Filtering and conversion of data into the different formats was performed using *Stacks, vcftools, PLINK* v.1.90 and *PGDspider* v.2.11.2. Datasets for creating site frequency spectra were filtered in a similar way, except no MAF cut-off was used in order to retain informative low frequency sites and a maximum of 10% missing data within each dataset (e.g. each ecotype pair) was allowed. For all analyses, only one SNP per locus was retained to reduce the effect of linkage.

### Summary statistics and analysis of population structure

To assess population structuring across and within lineages, and within lakes, we applied multiple approaches using the global and lineage-specific SNP datasets. First, we used PCA in *adegenet* (R-package) to assess major axes of genetic variation. Second, *Admixture* v.1.3 (Alexander & Novembre 2009) was used with a tenfold cross-validation to detect the most likely numbers of clusters and genetic ancestry proportions within lineages. *Genodive* v.2.0b27 (Meirmans & Tienderen 2004) was used to estimate pairwise genetic differentiation (Weir-Cockerham Fst) between ecotypes within and across lakes using the global dataset, with 10,000 permutations. We calculated genome-wide nucleotide diversity for each ecotype for all populations based on all SNPs using *vcftools*. We assessed the relationship among populations within and across lineages using a neighbour-joining splits network using *SplitsTree4 v.4.14.4* (Huson & Bryant 2005). To assess genetic co-ancestry among individuals within and across lakes, we used haplotype-based population inference approach implemented in *fineRADstructure* v.0.1 (Malinsky et al. 2018), using the same filtering criteria described for the SNP dataset. Analyses were performed using default settings.

### Introgression and differential admixture

We used *Treemix* v.1.13 (Pickrell & Pritchard 2012) to explore and visualize secondary gene flow within each lineage. We built maximum likelihood trees for non-admixed individuals (admixture threshold of 0.25 as inferred with *Admixture)* to reduce the effect of contemporary admixture. We fitted up to six and ten migration edges for the Atlantic and Siberian populations, respectively, and chose the most likely migration events based on the maximised variance explained, maximum likelihood and the significance of each migration edge. To formally test for introgression in a four-population tree (‘deviation from tree-ness’) we used *f4-statistics* implemented in the *fourpop* function of *Treemix.* Furthermore, to test for significant admixture within a population in a three-taxon comparison (Target taxon C; Reference taxon A, Reference taxon B) we used *f3-statistics* as implemented in the *threepop* function in *Treemix*.

### Inference of evolutionary histories

To distinguish between alternative evolutionary scenarios leading to ecotype diversity within lakes, we used coalescence simulations implemented in *fastsimcoal2* v.2.5.2.3 (Excoffier et al. 2013) and information contained in the multidimensional site frequency spectrum (SFS). 2-population and 3-population SFSs were created using *∂a∂i* v.1.6.3 (Gutenkunst et al. 2009)for each population and parapatric outgroups if appropriate. Populations were down-sampled by around 30% to reduce the effect of missing data. The minor folded site frequency spectrum was used due to the lack of a trinucleotide substitution matrix for salmonids and sequencing data for outgroup species. To determine absolute values for divergence times and other inferred parameters, we corrected the number of monomorphic sites in the SFS (Jacobs et al. 2018; Kautt et al. 2016). A mutation rate of 1×10^−8^ was used as no accurate mutation rate for Arctic charr or salmonids is available (Rougeux et al. 2017).

For all sympatric ecotype pairs, seven pairwise demographic models describing different historical divergence scenarios were tested (Supplementary Fig. S7): Strict isolation (SI), Ancient migration (AM), Isolation-with-migration (IM), Secondary contact (SC), Secondary Contact with introgression (AdmSC), Isolation-with-migration with a historical change in migration rate (IMchange) and an IM-model with a historical introgression event sympatric species (IMint) were tested. In lakes with three sympatric ecotypes (Kamkanda and Kalarskii Davatchan), or strong admixture across lakes (Loch Dughaill and Loch Uaine), models describing different combinations of strict isolation, isolation-with-migration, secondary contact, introgression and hybrid speciation were tested for all three ecotypes/populations together. These models tested in general two different evolutionary histories. The isolation-with-migration models test a history of divergence under constant gene flow (with differing rates), whereas secondary contact models mainly test the occurrence of historical secondary contact between different distinct lineages or populations (e.g. distinct gene pools or glacial refugial populations) prior to ecotype divergence. We ran a total of 30 iterations for each model and lake, and selected the most likely model based on the AIC (Excoffier et al. 2013). Each run consisted of 40 rounds of parameter estimation with 100,000 coalescent simulations. Point estimates of inferred parameters were taken from the most likely model and averaged over the top five runs.

### Patterns of selection and differentiation

To determine outlier loci between sympatric ecotypes, we screened the genome for loci showing genetic differentiation (Weir-Cockerham Fst calculated using *vftools)* above the 95^th^ quantile of the Fst distribution (outlier loci). We estimated the number and proportion of shared outlier SNPs and contigs containing outlier SNPs (but not the same SNPs across contigs) across benthivorous-planktivorous and piscivorous-planktivorous ecotype pairs. To determine if more outlier SNPs or contigs are shared among two ecotype pairs than expected by chance, we used *resample* (R-package) to resample n number of SNPs (n = number of outlier SNPs for each ecotype pair) 10,000 times with replacement from the full dataset and determined the mean number of shared SNPs from that distribution. We used a Chi-square test to determine if more outlier SNPs/contigs are shared than expected. We also plotted the standardised Fst (ZFst) against delta nucleotide diversity (Δ_π_ = π_ecotype2_ — π_ecotype1_) between sympatric ecotypes, to determine if outlier loci show reduced genetic diversity in one or both ecotypes, potentially indicating selective sweeps.

We used *pcadapt* v.3.0.4 (Luu et al. 2017) to identify SNPs under positive selection in each lineage. As the *pcadapt* approach is based on detecting loci that separate along different principal components, we used the first 10 and 14 principal component axes for the Atlantic and Siberian populations, respectively, as these separate the different lakes and sympatric ecotypes within lakes. We considered loci with *q-values* of 0.01 or below (false discovery rate of below 1%) to be candidates for being under positive selection.

### Genome-wide association analysis

To detect loci significantly associated with ecotype within each lineage we used a conditioned redundancy analysis (RDA), controlling for the effect of lake (condition) using *vegan* (R-package). Ecotypes were coded numerically, with planktivorous as 0, benthivorous as 1 and piscivorous as 2. We excluded unimodal populations and ecotypes from Tokko from this analysis. SNPs were selected as significantly associated with ecotype if the z-transformed loading for RDA1 (and RDA2 in the Siberian population) was above 2 or below −2 (equivalent to a two-tailed p-value < 0.05). To test if more ecotype-associated SNPs were shared across the lineages, we used the same resampling approach as for the outlier SNPs.

We imputed missing data in each SNP dataset using the *LD-kNNi* method implemented in *Tassel5* (Bradbury et al. 2007), based on the 10 closest genotypes using the default settings. To test the imputation accuracy, we calculated Pearson’s correlations between allele frequencies before and after imputation for the full dataset and the subset with the highest proportion of missing data.

### RNAseq and processing

Total RNA was extracted from white muscle tissue from 44 individuals (N=4 per ecotype per lake) from five lakes (Awe, Tay, Dug, Kam, Tok), representing all possible ecotypes (benthivorous, planktivorous, insectivorous, piscivorous), using PureLink RNA Mini kits (Life Technologies, Carlsbad, CA). Extractions were carried out following the manufacturer’s instructions, with the exception of an additional homogenisation step using a FastPrep-24 (MP Biomedicals), prior to isolation. RNA quantity and quality were assessed using the Qubit 2.0 fluorometer (Life Technologies, Carlsbad, CA) with HS Assay kits and a 2200 Tapestation (Agilent, Santa Clara, CA), respectively. High quality RNA was achieved, with A260/280 ratios between 1.9 and 2.1 and RNA Integrity Numbers above 8.3. RNA-seq libraries were prepared and sequenced at Glasgow Polyomics (University of Glasgow) for Awe, Tay, Dug and Kam and at BGI (Shenzhen, China) for Tok. Individual cDNA libraries were prepared for each individual using the TruSeq Stranded mRNA Sample Preparation kit (Illumina, San Diego, CA) in combination with a Poly-A selection step. Libraries were sequenced on an illumina NextSeq 500 (Illumina, San Diego, CA) using 75 bp paired-end sequencing, at a depth of 25-30M reads per library. Raw reads were processed using *Scythe v0.9944 BETA* (https://github.com/vsbuffalo/scythe/) and *Trimmomatic v0.36* (Bolger et al. 2014). Leading and trailing bases with a Phred quality score <20 were removed and a sliding window approach (4 bp window size) was used to trim reads at positions with Phred scores below 20. A minimum read length of 50 bp was allowed. We used *FastQC v0.11.2* to assess read-quality before and after processing. Processing removed ~2% of reads, resulting in 1.81 billion cleaned reads. The resulting reads were aligned against the Arctic charr draft genome using *STAR* v2.5.2b (Dobin et al. 2013), with default parameters. Raw reads were counted for each gene based on the longest isoform annotation using the HTSEQ-count python package (Anders et al. 2015) with the unstranded (--stranded=no), CDS—based (--type=CDS, --idattr=Parent) settings. Only genes with at least 20 read counts per lake were used for further downstream analyses.

### Gene expression analysis

To identify the major axes of expression variation across lakes, we performed a principal components analysis using the *svd* PCA approach in *pcaMethods* (R-epackage) based on rld-transformed gene expression data (transformed using the *DEseq2* R-package) (Love et al. 2014). The raw read count table was used for the differential gene expression analysis between sympatric ecotypes using *DESeq2* on a per lake basis. Furthermore, to identify genes with ecotype-associated expression patterns, we performed a conditioned RDA on rld-transformed count data using the *vegan* R-package controlling for lineage and lake (conditions). We again used z-transformed loadings above 2 or below −2 as the significance threshold.

To further examine the functional bases of trophic divergence Arctic charr, we used a Weighted Gene Co-Expression Network Analysis (WGCNA) to identify co-expressed gene modules (Langfelder & Horvath 2008). Network analyses were only performed on benthivorous—planktivorous ecotype pairs (from Awe, Tay, Dug and Kam; N=32, four per ecotype, per lake). To reduce stochastic background noise from lake-specific effects in our expression data, we used a linear mixed model in *variancePartition* (R-package) to identify a subset of genes with expression variation attributed to ecotype (see SI for detail). All genes for which ‘ecotype’ explained more than 10% of the total expression variation across individuals were used for network construction. A single network was constructed for all 32 samples and 1,512 ecotype-associated genes, from the log2 scaled count data (DESeq2: rlog), using *WGCNA* (R-package), following the standard procedure. Network modules were defined using the dynamic treecut algorithm, with a minimum module size of 25 genes and a cut height of 0.992. The module eigengene distance threshold was set to 0.25 to merge similar modules. To determine the significance of module-trait relationships, Pearson’s correlations were calculated between module eigengenes (the first principal component of the expression profile for a given module) and lake and ecotype. P-values were Benjamini-Hochberg corrected (FDR<0.05).

### *Cis*-eQTL mapping

To determine if the expression of candidate genes is genetically determined we performed *cis*-eQTL mapping using all benthic-pelagic ecotype pairs (N=32). First, we called SNPs from reference-aligned RNAseq data using *freebayes* (https://github.com/ekg/freebayyes), after marking duplicates using *picard*, with a coverage threshold of three. We only retained biallelic SNPs with a phred quality score above 30, a genotype quality above 20 and an allele-depth balance between 0.25 and 0.75. Furthermore, we filtered for Hardy-Weinberg disequilibrium (p-value threshold < 0.01), and only kept sites that were present in at least 90% of all individuals across populations. The filtering was performed using the *vcffilter* command implemented in *vflib* and *vftools.* Using these filtering steps, we retained 12,393 SNPs. To associated gene expression with sequence polymorphisms and identify *cis*-eQTL, we used *MatrixEQTL* v.2.2 (R-package) (Shabalin 2012). We used the linear model with lake and lineage as covariates, and a maximum distance of 1 Mbp between SNP and differentially expressed gene (cis-acting polymorphism only). Cis-eQTL were identified with a false-discovery rate (FDR) below 0.1 after correcting for multiple testing.

### Characterisation of differentially expressed genes

To detect genetic pathways associated with ecotype-associated differentially expressed genes (identified using RDA) and co-expressed gene modules, we performed overrepresentation analyses using the *WebGestalt* tool (Wang et al. 2017). We only used Arctic charr genes which had 1:1 or 2:1 orthologs in the zebrafish genome, with the following settings: minimum number of genes for a category=5, maximum number of genes for a category=500, number of permutations=1000, number of categories with leading-edge genes=20, KEGG pathways, organism=Danio *rerio*.

### Gene sharing between comparisons

To calculate the expected number of shared differentially expressed genes (DEGs) between comparisons we used a permutation-based approach, similar to the outlier comparison, with 10,000 permutations. We randomly sampled N genes (N=the number of DEGs in a comparison) 10,000 times from each dataset with replacement and calculated the expected number of shared DEGs as the mean number of shared resampled genes in each comparison. We calculated proportion-based p-values based on the number of observed and expected shared DEGs, and the total number of genes for each pairwise comparison using R function *prop.test.*

### Trajectory and regression analysis

Similar to the phenotypic trajectory analysis, we performed trajectory analyses (TA) based on different genetic datasets and gene expression data. The TA was performed using the *geomorph trajectory.analysis* function based on PC scores derived for each dataset. For the genetic data, we calculated trajectory lengths and angles for all neutral SNPs (N=7,179 SNPs, PC1-6; θ_Gn_, and ΔL_Gn_), outlier SNPs (N=986 SNPs, PC1-7; θ_Go_ and ΔL_Go_) and ecotype-associated SNPs (N=217 SNPs in the Atlantic lineage, PC1-4; N= 582 SNPs in the Siberian lineage, PC1-6; θ_RDA_, ΔL_RDA_). Except for the ecotype-associated SNPs, the TA was performed for both lineages combined. We also performed TA based on PC1-6 for all expressed genes (θ_GEx_, ΔL_GEx_) and for PC1-5 based on all genes associated with ecotype in the RDA (θ_canGEx_, ΔL_canGEx_). In all cases, we selected all PCs that cumulatively explain more than 50% of variation.

To identify how the different factors (phenotype, genotype, evolutionary history and gene expression) are correlated, we performed linear regression analyses using the *lm* function in R with different input datasets. First, we compared how differences in the angles between trajectories and lengths of trajectories, calculated based on different datasets (linear traits (θ_p_, ΔL_p_), neutral SNPs, outlier SNPs, ecotype-associated SNPs, overall gene expression and ecotype-associated expression), are correlated to identify if ecotype pair divergence one level explains the divergence patterns on a different level. Furthermore, we determined how absolute magnitudes of divergence (absolute length of trajectories for the same datasets (L), and neutral and outlier-based Fst) are correlated. We also tested the effect of ecosystem size on population nucleotide diversity and divergence patterns using linear regression analyses. We used the first PC from a PCA performed using *prcomp* based on maximum lake depth and surface area, estimated using Google Earth Pro, as a proxy for ecosystem size (Recknagel et al. 2017). Ecosystem size is used as a proxy for ecological opportunity within a system (Recknagel et al. 2017).

## Acknowledgements

N.V. Alekseyeva, R.S. Andreev, A.A. Kalinina, V.S. Khlystov, I.G. Khoro, I.B. Knizhin (deceased), A. Loshakov, A.N. Matveev, A.G. Osinov, M. Yu Pichugin, V.K. Pomazkin, V.V. Pulyarov, V.P. Samusenok, D.V. Schepotkin, F.N. Shkil’, A.A. Sokolov (deceased), S.D. Sviridov, A.I. Vokin, A.L. Yur’ev, I.I Yur’ev provided assistance with sampling, D.A. Akzhigitova, M. Yu Pichugin, A.A. Sokolov provided assistance with morphometric measures. We thank A. Adam for technical support in the lab, P.C Robinson for generating some sequence data, and Glasgow Polyomics for NGS.

## Authors contributions

KRE, CEA conceived the project; NVG, SSA, CEA, KRE designed experiments; NVG, SSA, MC, AJ, OH, CEA collected / contributed samples; BFK, JSL, EBR generated the draft genome scaffolds; AJ, MC generated data; AJ analyzed genomic and morphology data; MC, AY analyzed transcriptomic data; AJ, AY anchored and annotated the genome; KRE, CEA, AY supervised the project; AJ, KRE drafted the manuscript; all authors contributed to and approved the final manuscript.

## Data and material availability

Sequence data are available from [with acceptance].

## Author Information

Reprints and permissions information is available at www.nature.com/reprints. The authors declare no competing financial interests. Correspondence and requests for materials should be addressed to K.R.E.

## Supplementary Material

### Supplementary note: Genome annotation, orthology inference and chromosome assembly

We were able to assemble 62.6% of the draft genome (1.346 Gb) into 39 Arctic charr chromosomes with a total of 26.7% (0.573 Gb) being oriented. 37.4 % (0.804 Gb) of contigs could not be anchored to the linkage map. We annotated 44102 genes with 55443 alternative isoforms using homology-based *GeMoMa* gene prediction software. The BUSCO analysis estimated that the annotation contained 88% of completely assembled BUSCOs and 3.4% of fragmented BUSCOs, while 7.6% of BUSCOs were missing. Among the 88% completely assembled BUSCOs, 62% were duplicated, which is consistent with the estimation for the salmon genome. Orthofinder returned 15485 orthogroups between Arctic charr and zebrafish with a mean and median orthogroup size equal to 2.7 and 2, respectively, which can be explained by the relatively recent whole genome duplication in salmonids. We detected a total of 8428 single-copy orthogoups between Arctic charr and zebrafish.

### Supplementary Figures

**Supplementary Fig S1.**
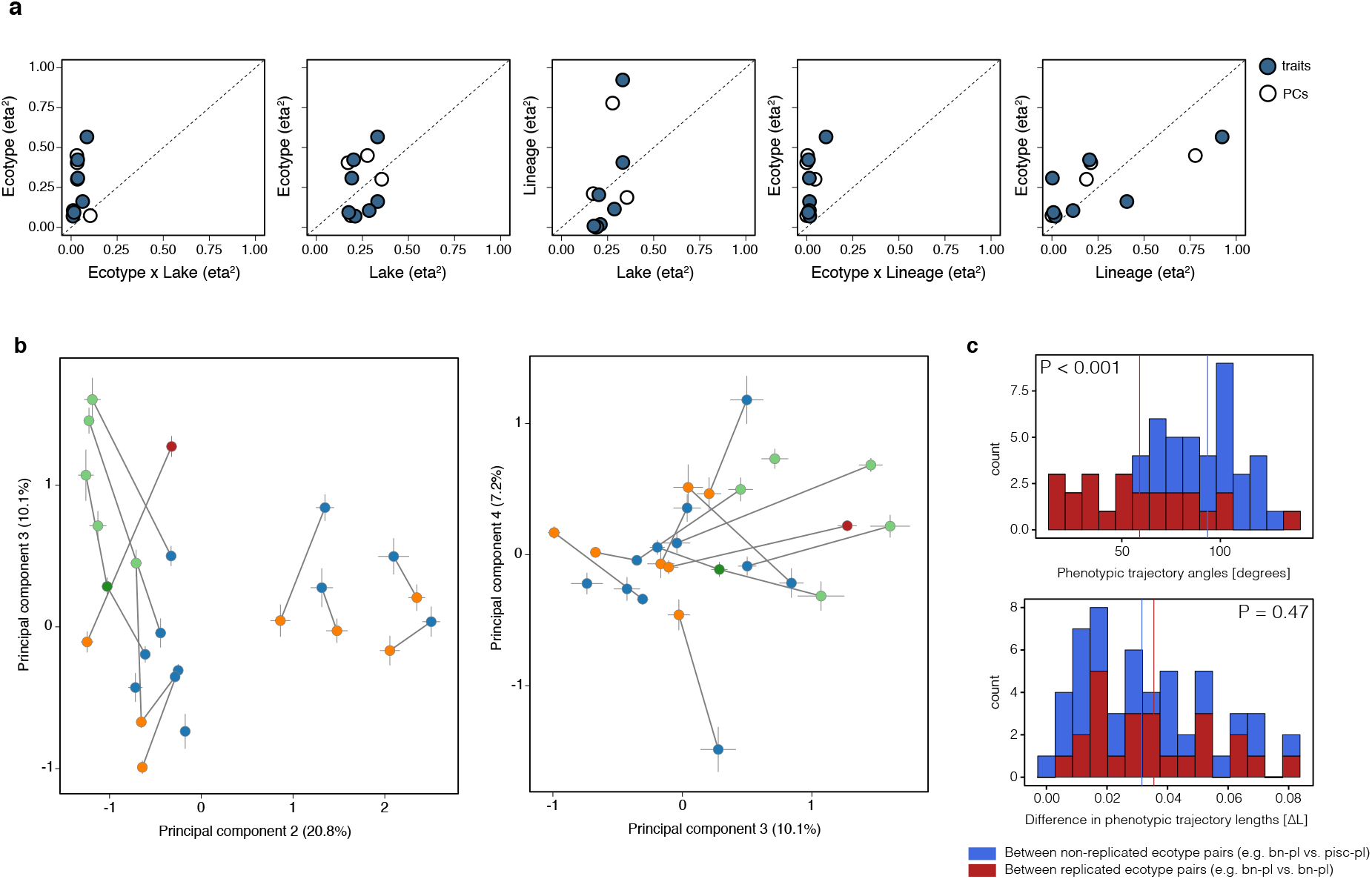
Phenotypic parallelism. **(a)** Effect sizes (partial η^2^) of linear model terms for each phenotypic trait and PC1 to PC4 of the linear trait principal component analysis. **(b)** Principal component plots for PC1 vs PC2 and PC3 vs PC4, with points showing the centroids and standard error for each ecotype and sympatric ecotype pairs are connected by lines. Points are coloured by ecotype: blue – planktivorous, orange – benthivorous, green – piscivorous, and red – insectivorous. **(c)** Distribution of phenotypic trajectory angles and differences in phenotypic trajectory lengths for comparisons between replicated ecotype-pairs (N=24) and between non-replicated ecotype pairs (N=30). The means for each dataset are shown by the solid lines and the p-value shows the result of a Wilcoxon rank sum test testing the difference in the mean between both datasets.

**Supplementary Fig S2.**
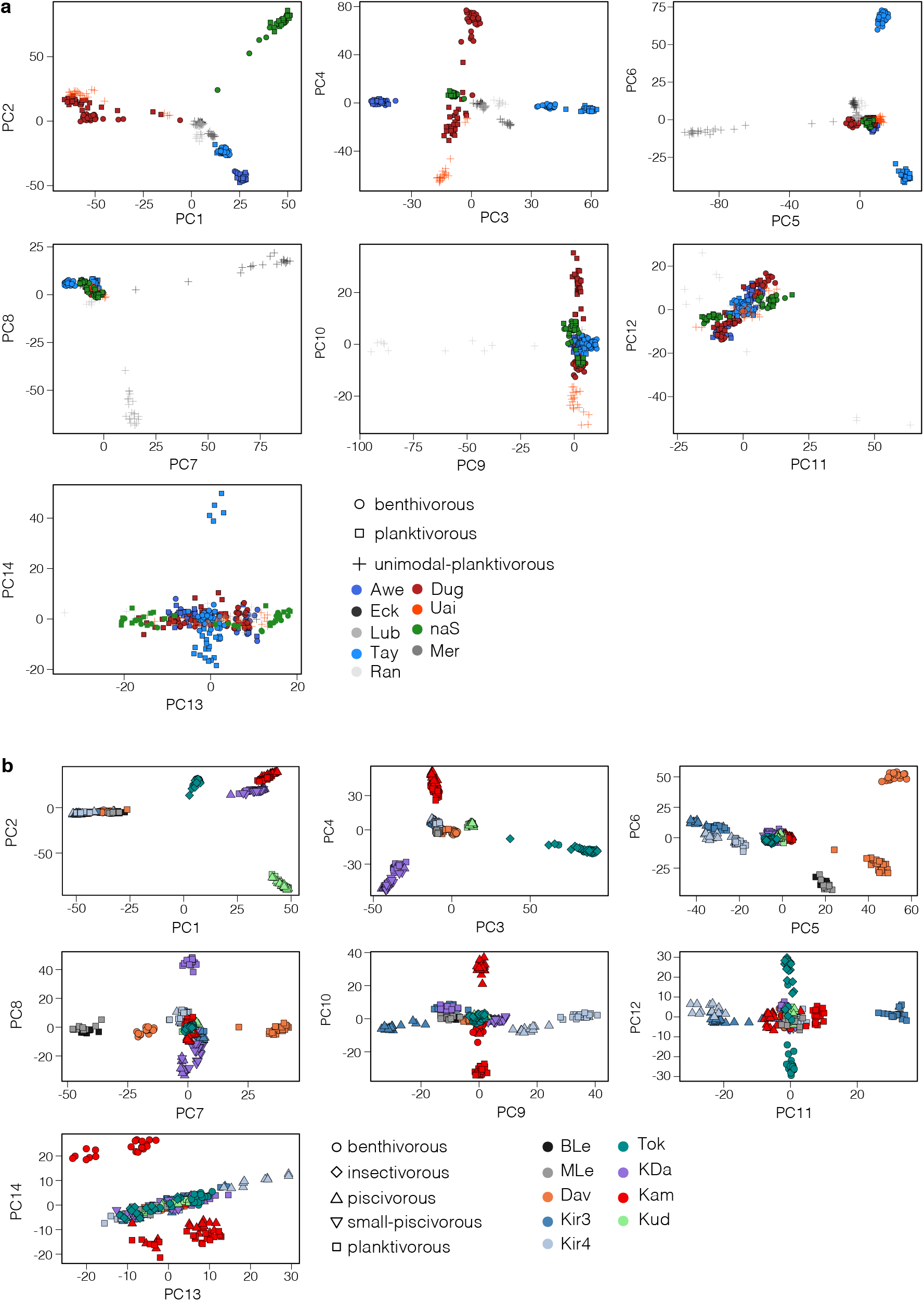
Principal component plots by lineage. **(a)** Principal component plots based on 6,039 SNPs showing PC1 to PC14 for individuals from the Atlantic lineage (N=300 ind.). **(b)** Principal component plots for all individuals from the Siberian lineage (N=328 ind.) based on 4,475 SNPs. Individual points are shaped based on ecotype and coloured by lake.

**Supplementary Fig S3.**
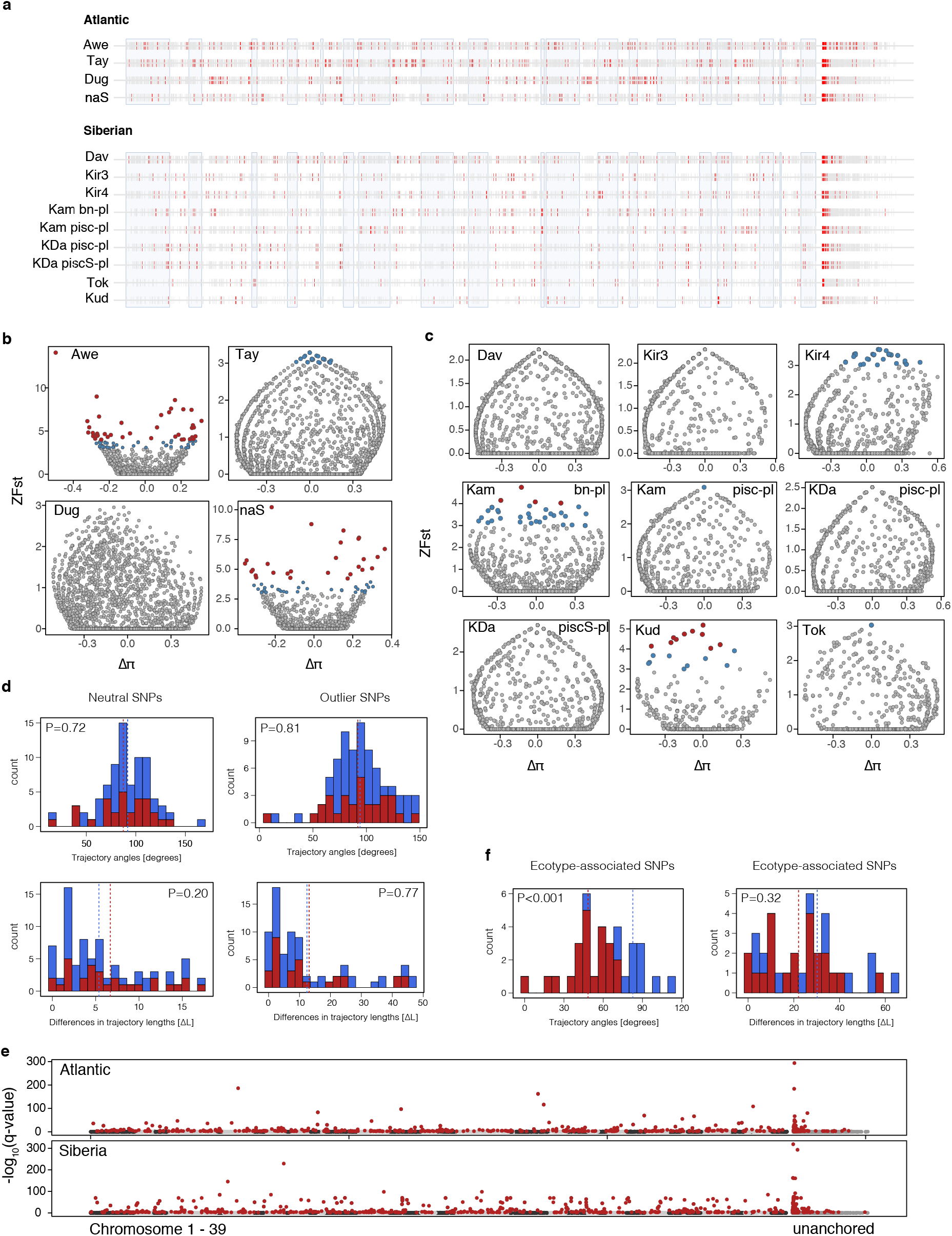
Genetic differentiation and selection. **(a)** Genome scan plot with top 5%-quantile Fst outlier loci highlighted in red. Chromosomes are alternatingly highlighted by blue boxes, and unplaced scaffolds are placed on the right side. **(b,c)** Genome-wide patterns of differentiation between sympatric ecotypes in the **(b)** Atlantic and **(c)** Siberian lineage. The z-transformed Fst (ZFst) is plotted against the delta nucleotide-diversity (Δπ; bn – pl and pisc – pl). If the Δπ deviates from zero, it shows genetic diversity at this locus is reduced in one of the ecotypes. Loci with ZFst values above 4 were inferred to be significantly differentiated (red dots) and loci with ZFst above 3 (blue dots) are also reported. **(d)** Distribution of trajectory angles and differences in trajectory lengths based on neutral SNPs (N=7,179) and outlier SNPs (N=986) for comparisons between replicated ecotype-pairs (N=30) and between non-replicated ecotype pairs (N=48). The means for both datasets are shown as dotted lines and the p-value shows the result of a Wilcoxon rank sum test testing the difference in the mean between both datasets. **(e)** Manhattan plots showing the *pcadapt* results (-log10[q-value]) across the Arctic charr genome. Loci with significant signals of differentiation and selection are highlighted in red. **(f)** Distribution of trajectory angles and differences in trajectory lengths based on ecotype-associated SNPs in the Atlantic (N=217) and Siberian lineage (N=582) for comparisons between replicated ecotype-pairs (N=22) and between non-replicated ecotype pairs (N=12). The mean for each dataset is shown as a dotted line and the p-value shows the result of a Wilcoxon rank sum test testing the difference in the mean between both datasets.

**Supplementary Fig S4.**
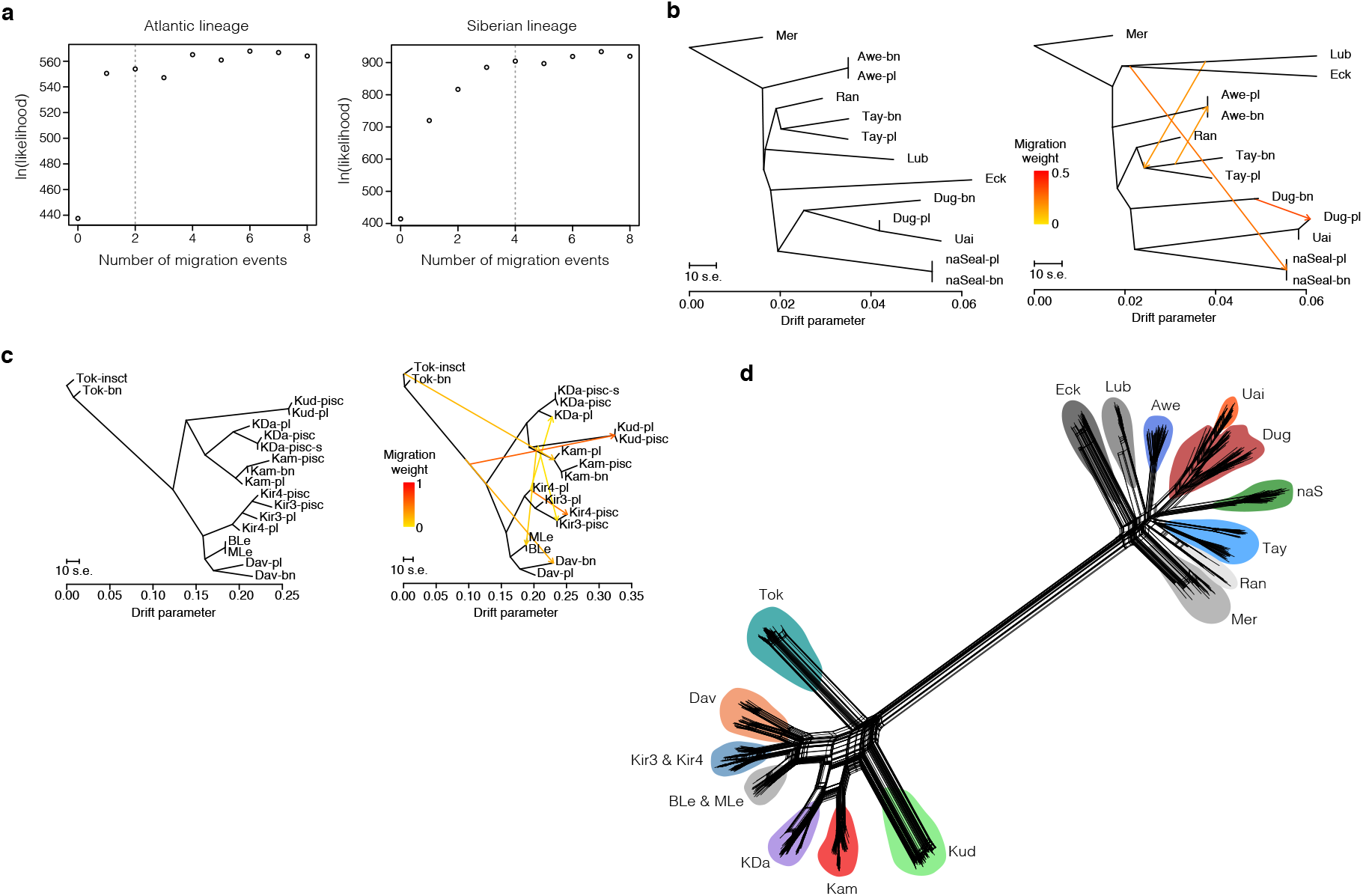
Treemix maximum-likelihood (ML) trees and Splitstree network. **(a)** Likelihood of trees with different numbers of fitted migration events for the Atlantic and Siberian lineage. **(b)** ML-trees from *Treemix* with zero and four migration edges respectively for all populations from the Atlantic lineage. Migration edges are shown as arrows coloured by migration weight. **(c)** *Treemix* ML-trees for all populations from the Siberian lineage with zero and six migration events fitted. **(d)** Phylogenetic Splitstree network for all individuals (N=630) from the Atlantic and Siberian lineage. Population codes are described in Supplementary table S1.

**Supplementary Fig S5.**
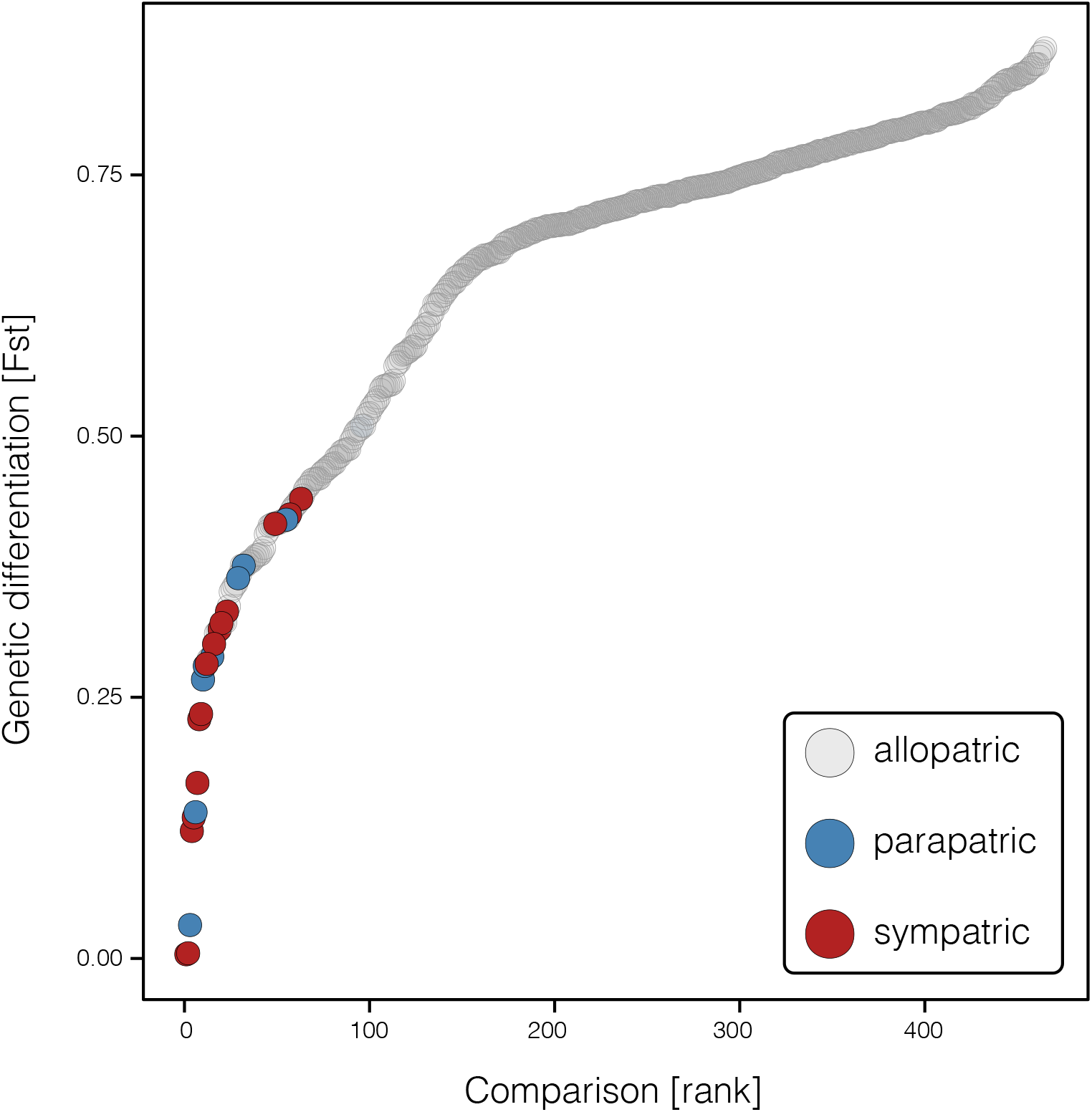
Continuum of genetic differentiation. The degree of pairwise genetic differentiation (Fst) calculated based on 12,215 SNPs is plotted against its rank. Points are coded based on comparison; grey = allopatric comparison between ecotypes from different lakes, blue = parapatric comparison between ecotypes from adjacent connected lakes from the same catchment, sympatric comparison between ecotypes from the same lake. Note that some sympatric ecotype pairs show higher degrees of genetic differentiation than ecotypes in allopatric comparisons.

**Supplementary Fig S6.**
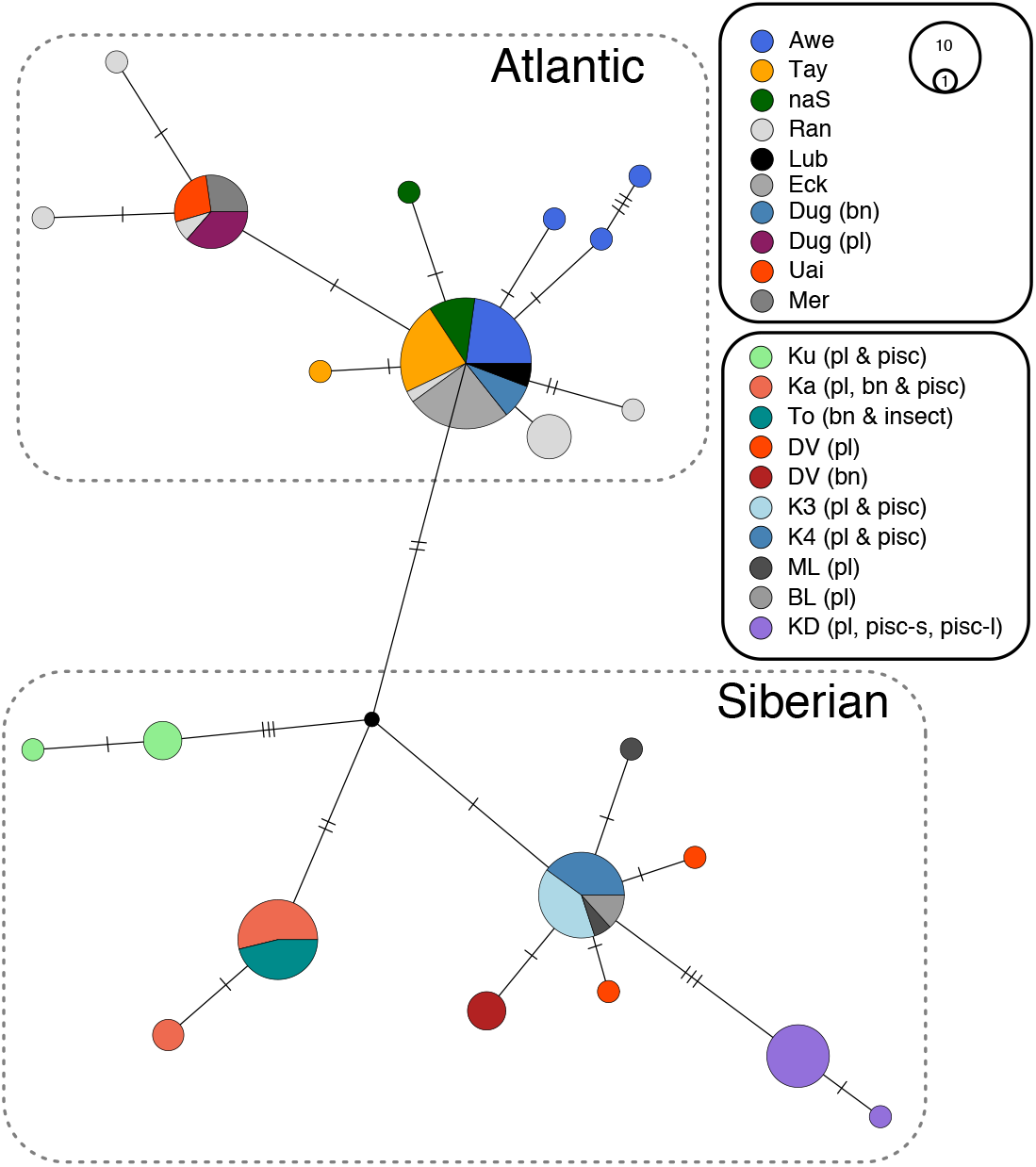
Haplotype network for the mitochondrial *ND1* gene (N=107). The size of each circle corresponds to the number of individuals sharing a haplotype. When sympatric ecotypes share one or several haplotypes than the circles or pies are only coloured by lake of origin. However, when sympatric ecotypes have distinct haplotypes, then each circle or pie is coloured by ecotype.

**Supplementary Fig S7.**
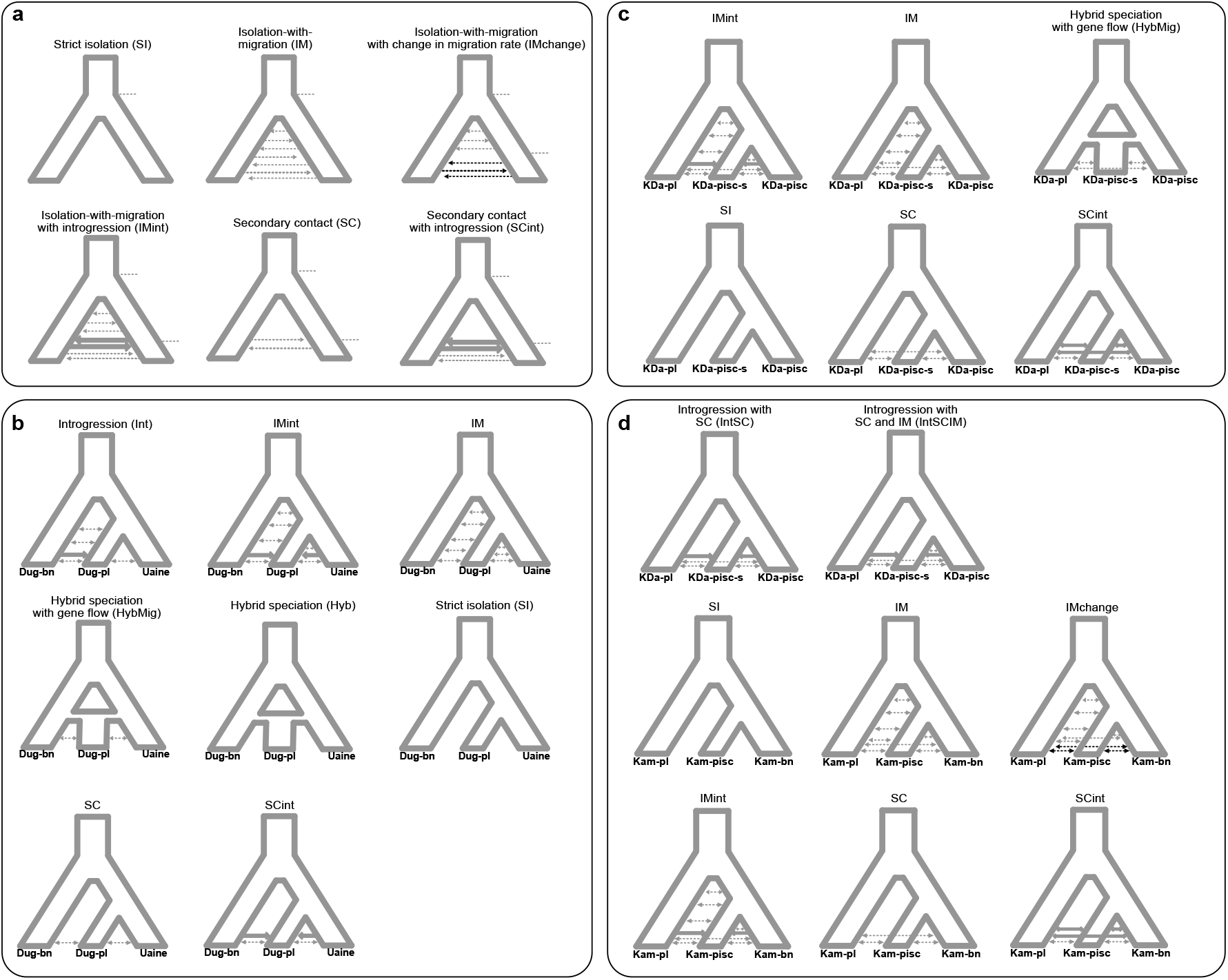
Overview of all tested demographic models in *fastsimcoal2.* **(a)** Illustrations of two-population models tested. **(b)** Three-population models tested for the evolutionary history of Dughaill and Uaine. **(c)** All three-population models tested for Kalarskii-Davatchan and **(d)** Kamkanda. We inferred parameters, such as divergence times, timing of secondary contact and introgression, strength of introgression and gene flow and effective population sizes.

**Supplementary Fig S8.**
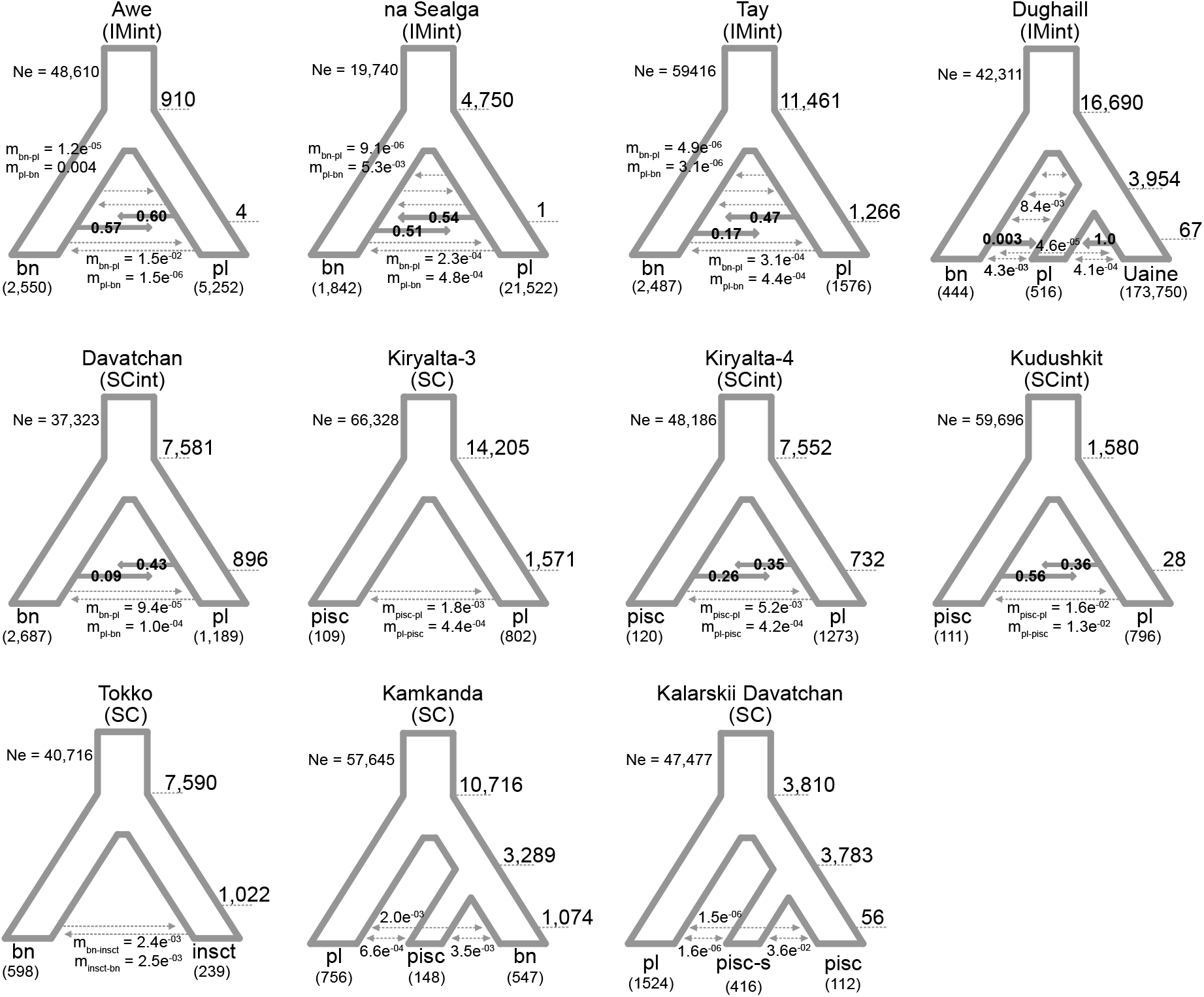
Illustrations of the most likely evolutionary history for each population, and inferred parameters using *fastsimcoal2.* All parameters are point estimates that were averaged across the five runs with the highest likelihood.

**Supplementary Fig S9.**
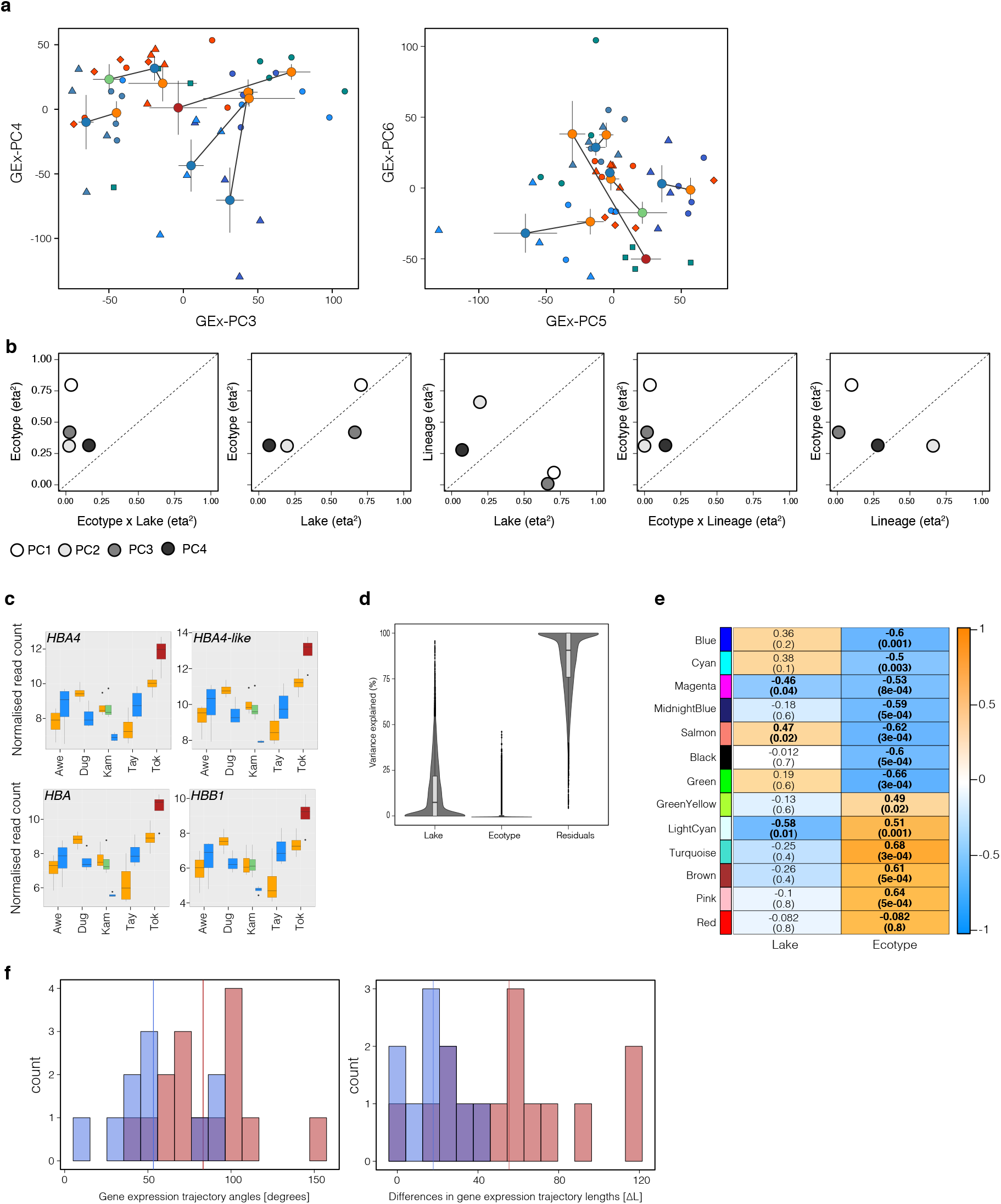
Gene expression results. **(a)** Principal component (PCA) plots based on gene expression data for PC3 vs PC4 and PC5 vs PC6. Individuals are shown by individual points shaped by ecotype and coloured by lake of origin. Centroids for each ecotype are shown including standard error and coloured by ecotype (blue – planktivorous, orange – benthivorous, green – piscivorous, red – insectivorous). Centroids of sympatric ecotypes are connected by a line. **(b)** Liner model term effect sizes (partial h^2^) for PC1 to PC4 from the gene expression PCA. **(c)** Boxplots showing the normalised expression of different haemoglobin paralogs across ecotypes and lakes. **(d)** Distribution of explained variances for each transcript by model term for the linear mixed effects model of gene expression. **(e)** Correlation between expression of WGCNA module eigengenes and lake (population of origin) and ecotype. Each row represents a module of co-expressed genes (identified by colour). Spearman’s correlation coefficients for the module expression –variable correlations are given in each cell. The corresponding corrected p-values are given in parenthesis. Significant correlations are highlighted in bold. Cells are coloured based on their correlation, with orange cells being correlated with up regulation of gene expression in benthivorous ecotypes and blue being associated with up-regulation in planktivorous ecotypes. **(f)** Distribution of gene expression trajectory angles and differences in trajectory lengths for within and between ecotype comparisons. Comparisons are coloured by comparisons between replicated ecotype-pairs (red; N=6) and between non-replicated ecotype pairs (blue; N=9). The means for each dataset are shown by the solid lines. Means do not differ between the comparisons (P > 0.05).

**Supplementary Table S1.** Sampling sites and sample sizes for populations used for phenotypic and genomic analysis of Arctic charr from the Atlantic and Siberian lineage.

| Atlantic |  |  |  |  |  |  |  |  |  |  |  |  |
| --- | --- | --- | --- | --- | --- | --- | --- | --- | --- | --- | --- | --- |
| Lake | Abb<sup>1</sup> | Catchment | Lat. (N) | Lon. (W) | N phenotyped individuals |  |  | N genotyped individuals |  |  |  |  |
|  |  |  |  |  | Bn | Pl | Uni | Bn | Pl | Uni |  |  |
| Tay | Tay | Tay | 56°30' | 004°10' | 43 | 76 | - | 21 | 31 | - |  |  |
| Rannoch | Ran | Tay | 56°41' | 004°16' | - | - | - | - | - | 11 |  |  |
| Awe | Awe | Awe | 56°20' | 005°05' | 34 | 36 | - | 25 | 28 | - |  |  |
| Eck | Eck | Cowal | 56°01' | 004°58' | - | - | - | - | - | 19 |  |  |
| Lubnaig | Lub | Forth | 56°17' | 004°17' | - | - | - | - | - | 20 |  |  |
| Dughaill | Dug | Ewe | 57°28' | 005°20' | 44 | 52 | - | 31 | 31 | - |  |  |
| Uaine | Uai | Ewe | 57°31' | 005°23' | - | - | - | - | - | 23 |  |  |
| naSealga | naS | - | 57°79' | 004°30' | 41 | 19 | - | 19 | 21 | - |  |  |
| Merkland | Mer | Shin | 58°14' | 004°44' | - | - | - | - | - | 22 |  |  |
| Siberian |  |  |  |  |  |  |  |  |  |  |  |  |
| Lake | Abb<sup>1</sup> | Catchment | Lat. (N) | Lon. (E) | N phenotyped individuals |  |  |  | N genotyped individuals |  |  |  |
|  |  |  |  |  | Bn | Pl | Pisc | Insct | Bn | Pl | Pisc | Insct |
| Kudushkit | Kud | Chaya | 56°33' | 110°37' | - | - | 20 | - | - | 20 | 19 | - |
| Davatchan | Dav | Chara, Olekma | 56°27' | 117°33' | 76 | 114 | - | - | 20 | 25 | - | - |
| Kalarskii Davatchan | KDa | Kalar,Vitim | 56°16.5' | 116°38.5' | - | 83 | 15/81<sup>2</sup> | - | - | 19 | 12/21<sup>2</sup> | - |
| Kiryalta-3 | Kir3 | Chara, Olekma | 57°08.5' | 119°27' | - | 23 | 30 | - | - | 20 | 19 | - |
| Kiryalta-4 | Kir4 | Chara, Olekma | 57°06.5' | 119°28' | - | 42 | 25 | - | 5 | 21 | 13 | - |
| Tokko | Tok | Tokko, Chara, Olekma | 57°11' | 119°41' | 42 | - | - | 47 | 18 | - | - | 19 |
| Kamkanda | Kam | Khani, Olekma | 57°05.5' | 199°48.5' | 146 | 163 | 26 | - | 20 | 19 | 20 | - |
| Bol'shoe Leprindo | BLe | Chara, Olekma | 56°37' | 117°31' | 25 | - | - | - | 10 | - | - | - |
| Maloe Leprindo | MLe | Chara, Olekma | 56°36' | 117°22' | 26 | - | - | - | 11 | - | - | - |

Note: <sup>1</sup>Abbreviations used for these populations in the manuscript. <sup>2</sup> There are two piscivorous ecotypes in Kalarskii Davatchan, large piscivorous / small piscivorous ecotype. Ecotype abbreviations: Bn – benthivorous, PL – planktivorous, Uni – unimodal-planktivorous populations or without phenotypic information (Rannoch), Pisc – piscivorous, Insct – insectivorous.

**Supplementary Table S2.** Effect sizes (partial h^2^) for each model term from trait-by-trait linear models.

| Trait/PC | Ecotype ($\eta^2$) | Ecotype x Lake ($\eta^2$) | Lake ($\eta^2$) | Lineage ($\eta^2$) | Ecotype x Lineage ($\eta^2$) |
| --- | --- | --- | --- | --- | --- |
| **phen.PC1** | 0.3012 | 0.0338 | 0.355 | 0.1873 | 0.0452 |
| **phen.PC2** | 0.4499 | 0.031 | 0.2772 | 0.7783 | 0.0046 |
| **phen.PC3** | 0.405 | 0.0329 | 0.172 | 0.211 | 0.0000 |
| **phen.PC4** | 0.0732 | 0.1039 | 0.1874 | 0.0001 | 0.0001 |
| **HDO** | 0.5666 | 0.0857 | 0.332 | 0.9235 | 0.1048 |
| **HDE** | 0.1616 | 0.0624 | 0.3327 | 0.4066 | 0.0152 |
| **HL** | 0.0701 | 0.0102 | 0.2108 | 0.018 | 0.0166 |
| **PFL** | 0.4232 | 0.0356 | 0.2024 | 0.2041 | 0.0094 |
| **ED** | 0.3083 | 0.0360 | 0.1923 | 0.0025 | 0.0144 |
| **ML** | 0.1051 | 0.0114 | 0.287 | 0.1143 | 0.0146 |
| **LJL** | 0.0932 | 0.0159 | 0.1758 | 0.0083 | 0.0086 |

Note: phen.PC – PC1 to PC4 from principal components analysis performed on all seven linear traits. HDO – head depth at operculum, HDE – head depth at eye, HL – head length, PFL – pectoral fin length, ED – eye diameter, ML – maxilla length, LJL – lower jaw length.

**Supplementary Table S3.** Results of phenotypic trajectory analysis based on all seven linear traits. Differences in trajectory lengths (ΔL, upper part) or angles (0, lower part) below the diagonal and p-values are above the diagonal.

| $\Delta L$/p-value | Awe_bn-pl | Dav_bn-pl | Dug_bn-pl | Kam_bn-pl | Kam_pl-pisc | KDa_pl-pisc-s | KDa_pl-pisc | Kir3_pisc-pl | Kir4_pisc-pl | naS_bn-pl | Tay_bn-pl |
| --- | --- | --- | --- | --- | --- | --- | --- | --- | --- | --- | --- |
| Awe_bn-pl | 0 | 0.001 | 0.04 | 0.001 | 0.078 | 0.683 | 0.001 | 0.002 | 0.04 | 0.584 | 0.254 |
| Dav_bn-pl | 0.0699 | 0 | 0.013 | 0.049 | 0.01 | 0.001 | 0.512 | 0.979 | 0.061 | 0.001 | 0.001 |
| Dug_bn-pl | 0.0347 | 0.0352 | 0 | 0.244 | 0.824 | 0.037 | 0.185 | 0.056 | 0.811 | 0.017 | 0.336 |
| Kam_bn-pl | 0.0499 | 0.0200 | 0.0152 | 0 | 0.143 | 0.001 | 0.507 | 0.195 | 0.485 | 0.003 | 0.02 |
| Kam_pl-pisc | 0.0311 | 0.0388 | 0.0036 | 0.0188 | 0 | 0.083 | 0.123 | 0.044 | 0.625 | 0.033 | 0.49 |
| KDa_pl-pisc-s | 0.0061 | 0.0638 | 0.0285 | 0.0437 | 0.0249 | 0 | 0.003 | 0.001 | 0.036 | 0.291 | 0.322 |
| KDa_pl-pisc | 0.0594 | 0.0105 | 0.0247 | 0.0095 | 0.0283 | 0.0532 | 0 | 0.598 | 0.311 | 0.002 | 0.023 |
| Kir3_pisc-pl | 0.0703 | 0.0004 | 0.0356 | 0.0204 | 0.0392 | 0.0641 | 0.0109 | 0 | 0.139 | 0.001 | 0.009 |
| Kir4_pisc-pl | 0.0395 | 0.0304 | 0.0049 | 0.0103 | 0.0085 | 0.0334 | 0.0198 | 0.0307 | 0 | 0.015 | 0.255 |
| naS_bn-pl | 0.0110 | 0.0809 | 0.0456 | 0.0608 | 0.0420 | 0.0171 | 0.0703 | 0.0812 | 0.0505 | 0 | 0.096 |
| Tay_bn-pl | 0.0197 | 0.0502 | 0.0150 | 0.0302 | 0.0114 | 0.0135 | 0.0397 | 0.0506 | 0.0199 | 0.0307 | 0 |
| $\theta$/p-value | | | | | | | | | | | |
| Awe_bn-pl | 0 | 0.001 | 0.001 | 0.001 | 0.001 | 0.001 | 0.001 | 0.001 | 0.001 | 0.001 | 0.005 |
| Dav_bn-pl | 73.04 | 0 | 0.02 | 0.271 | 0.001 | 0.001 | 0.001 | 0.001 | 0.001 | 0.001 | 0.001 |
| Dug_bn-pl | 100.05 | 32.57 | 0 | 0.066 | 0.001 | 0.001 | 0.002 | 0.001 | 0.001 | 0.003 | 0.001 |
| Kam_bn-pl | 85.81 | 14.20 | 24.40 | 0 | 0.001 | 0.001 | 0.002 | 0.001 | 0.001 | 0.001 | 0.001 |
| Kam_pl-pisc | 87.22 | 120.36 | 129.13 | 119.15 | 0 | 0.001 | 0.001 | 0.742 | 0.007 | 0.001 | 0.001 |
| KDa_pl-pisc-s | 109.75 | 67.19 | 67.91 | 57.54 | 72.49 | 0 | 0.503 | 0.003 | 0.05 | 0.001 | 0.001 |
| KDa_pl-pisc | 115.10 | 74.48 | 68.77 | 63.23 | 69.84 | 17.87 | 0 | 0.022 | 0.35 | 0.001 | 0.001 |
| Kir3_pisc-pl | 94.41 | 111.25 | 115.13 | 107.00 | 16.46 | 58.83 | 54.08 | 0 | 0.126 | 0.001 | 0.001 |
| Kir4_pisc-pl | 98.00 | 78.35 | 78.59 | 71.50 | 52.45 | 33.51 | 26.81 | 37.22 | 0 | 0.001 | 0.001 |
| naS_bn-pl | 133.48 | 104.88 | 86.79 | 97.24 | 101.70 | 92.05 | 88.51 | 101.18 | 103.43 | 0 | 0.001 |
| Tay_bn-pl | 48.08 | 45.61 | 71.44 | 57.55 | 102.15 | 86.44 | 99.48 | 104.24 | 91.10 | 121.13 | 0 |

**Supplementary Table S4.** Results of Fst genome scans between sympatric ecotypes.

| Lake | Eco | mean Fst | outlier Fst | 95<sup>th</sup> perc. | N. outlier | N.ZFst>2 | N.ZFst>3 | N.ZFst>4 | N. fixed SNPs | % fixed |
| --- | --- | --- | --- | --- | --- | --- | --- | --- | --- | --- |
| **Awe** | Bn-Pl | 0.0113 | 0.1012 | 0.0653 | 175 | 185 | 86 | 40 | 0 | 0.00 |
| **Tay** | Bn-Pl | 0.2262 | 0.8483 | 0.7391 | 209 | 273 | 36 | 0 | 4 | 0.10 |
| **Dughail** | Bn-Pl | 0.2524 | 0.7537 | 0.6814 | 178 | 142 | 0 | 0 | 0 | 0.00 |
| **naSealga** | Bn-Pl | 0.0136 | 0.1195 | 0.0795 | 131 | 146 | 60 | 27 | 0 | 0.00 |
| **Davatchan** | Bn-Pl | 0.3291 | 0.9898 | 0.9523 | 113 | 127 | 0 | 0 | 76 | 3.38 |
| **Kamkanda** | Bn-Pl | 0.1336 | 0.6300 | 0.4935 | 96 | 99 | 36 | 4 | 0 | 0.00 |
| **Kamkanda** | Pl-Pisc | 0.2445 | 0.8567 | 0.7592 | 90 | 106 | 1 | 0 | 1 | 0.06 |
| **Kalarskii Dv** | Pl-Pisc | 0.2976 | 0.9569 | 0.8748 | 85 | 99 | 0 | 0 | 35 | 2.08 |
| **Kalarskii Dv** | Pl-Pisc-s | 0.2587 | 0.9393 | 0.8549 | 83 | 102 | 0 | 0 | 18 | 1.08 |
| **Kiryalta-3** | Pl-Pisc | 0.3074 | 0.9530 | 0.8845 | 59 | 51 | 0 | 0 | 15 | 1.26 |
| **Kiryalta-4** | Pl-Pisc | 0.1837 | 0.8208 | 0.6731 | 86 | 112 | 36 | 0 | 0 | 0.00 |
| **Kudushkit** | Pl-Pisc | 0.1003 | 0.6621 | 0.4677 | 37 | 40 | 19 | 10 | 1 | 0.14 |
| **Tokko** | Bn-Insect | 0.2233 | 0.8593 | 0.7557 | 46 | 54 | 2 | 0 | 2 | 0.22 |

Note: Bn – benthivorous, Pl – planktivorous, Pisc – piscivorous, Pisc-s – small-piscivorous, Insect – insectivorous. 95<sup>th</sup> perc. – 95<sup>th</sup> percentile of Fst distribution used for identifying outliers. N.outlier – Number of outlier loci identified. N.ZFst – number of SNPs with a z-transformed Fst above the threshold (2, 3 or 4). N.fixed SNPs – Number of fixed SNPs between ecotypes. % fixed – Percentage of fixed SNPs compared to all SNPs in the comparison.

**Supplementary Table S5.** Genes containing ecotype-associated SNPs identified in the cRDA analysis in both lineages.

| Contig | SNP | Position | Gene (within/downstream) | Gene product | Gene (upstream) | Gene product |
| --- | --- | --- | --- | --- | --- | --- |
| **Contig1179** | 5257_34 | 2295862 | downstream of RNA29111_R0 | receptor-type tyrosine-protein phosphatase eta |  |  |
| **Contig1218** | 6352_25 | 1446318 | RNA27331_R1 | sodium/potassium-transporting ATPase subunit beta-1-interacting protein 2-like |  |  |
| **Contig1811** | 20016_37 | 380212 | RNA101202_R1 | neuropilin-1a-like |  |  |
| **Contig2079** | 23905_27 | 770770 | RNA43764_R0 | zinc finger protein 536 | RNA43780_R0 | zinc finger protein 507-like |
| **Contig4128** | 42487_17 | 1064984 | RNA58831_R0 | putative ferric-chelate reductase 1 | RNA58840_R0 | calsequestrin-1 |
| **Contig4165** | 42701_25 | 784583 | RNA73507_R2 | sericin-1-like | RNA73498_R0 | pleckstrin |
| **Contig826** | 60249_16 | 1161922 | downstream of RNA76031_R1 | uncharacterized protein K02A2.6-like |  |  |
| **Contig949** | 65145_41 | 239650 | RNA98260_R0 | RNA polymerase II subunit A C-terminal domain phosphatase |  |  |

**Supplementary Table S6.** Results of the*f4-* and*f3-*-statistics for both lineages.

| **$f_4$-statistics** |  |  |  |  |  |  |
| --- | --- | --- | --- | --- | --- | --- |
| **PopA** | **PopB** | **PopC** | **PopD** | **$f_4$** | **se** | **Z-score** |
| **Dav-bn** | BLe | KDa-pisc | KDa-pisc-s | -0.00267318 | 0.00055094 | -4.85202 |
| **Dav-bn** | BLe | Kud-pisc | Kud-pl | 0.00172862 | 0.000546439 | 3.16342 |
| **Dav-bn** | Dav-pl | KDa-pisc | KDa-pl | -0.00281651 | 0.00142346 | -1.97865 |
| **Kam-pl** | Kam-bn | KDa-pisc | KDa-pisc-s | -0.00056964 | 0.000272173 | -2.09294 |
| **Kam-pl** | Kam-pisc | KDa-pisc | KDa-pisc-s | -0.00135753 | 0.000374307 | -3.62679 |
| **KDa-pisc** | KDa-pisc-s | Tok-bn | Tok-insct | -0.000578487 | 0.000183728 | -3.14862 |
| **KDa-pisc** | KDa-pisc-s | MLe | BLe | -0.000742263 | 0.000269313 | -2.75613 |
| **Kir4-pisc** | Kir4-pl | Kud-pisc | Kud-pl | -0.000837066 | 0.000326568 | -2.56322 |
| **Kir4-pisc** | Kir4-pl | KDa-pisc | KDa-pisc-s | 0.000539178 | 0.00026962 | 1.99977 |
| **Kud-pisc** | Kud-pl | KDa-pisc | KDa-pisc-s | -0.00175827 | 0.000372013 | -4.72636 |
| **Kud-pisc** | Kud-pl | KDa-pisc | KDa-pl | -0.00174521 | 0.000525462 | -3.32129 |
| **Kud-pisc** | Kud-pl | MLe | BLe | 0.000717697 | 0.000270048 | 2.65766 |
| **Dug-bn** | Eck | Dug-pl | Uai | 0.0216789 | 0.00161344 | 13.4364 |
| **Dug-bn** | Lub | Dug-pl | Uai | 0.0216151 | 0.00153642 | 14.0685 |
| **Tay-pl** | Tay-bn | Awe-pl | Dug-pl | -0.00484982 | 0.00211385 | -2.29431 |
| **Tay-pl** | Tay-bn | Awe-pl | Eck | -0.00888889 | 0.00248355 | -3.5791 |
| **Tay-pl** | Tay-bn | Awe-pl | Mer | -0.00576767 | 0.00231408 | -2.49242 |
| **Tay-pl** | Tay-bn | Awe-pl | naSeal-pl | -0.00527456 | 0.00239611 | -2.2013 |
| **Tay-pl** | Tay-bn | Awe-pl | Uai | -0.00525318 | 0.00238727 | -2.2005 |
| **$f_3$-statistics** |  |  |  |  |  |  |
| **Focal** | **Ref1** | **Ref2** | **$f_3$** | **se** | **Z-score** |  |
| **Dug-pl** | Dug-bn | Uai | -0.0259701 | 0.00110819 | -23.4348 |  |
| **Dug-pl** | Tay-bn | Uai | -0.00475052 | 0.00127475 | -3.72663 |  |
| **Dug-pl** | Ran | Uai | -0.00448714 | 0.00123844 | -3.62322 |  |
| **Dug-pl** | Tay-pl | Uai | -0.00434716 | 0.00126704 | -3.43097 |  |
| **Dug-pl** | Uai | Lub | -0.00435496 | 0.00140769 | -3.0937 |  |
| **Dug-pl** | Awe-pl | Uai | -0.0037364 | 0.00131955 | -2.83157 |  |
| **Dug-pl** | Eck | Uai | -0.0042912 | 0.00153863 | -2.78897 |  |
| **Dug-pl** | Uai | Awe-bn | -0.00371748 | 0.00133324 | -2.7883 |  |
| **Dug-pl** | Uai | naSeal-pl | -0.00329079 | 0.00140305 | -2.34546 |  |
| **Dug-pl** | naSeal-bn | Uai | -0.00295874 | 0.00140562 | -2.10494 |  |
| **Dug-pl** | Uai | Mer | -0.0030708 | 0.00146056 | -2.10249 |  |
| **KDa-pisc-s** | KDa-pisc | KDa-pl | -0.00249944 | 0.000420217 | -5.94796 |  |
| **KDa-pisc-s** | Kud-pisc | KDa-pl | -0.00191366 | 0.000606432 | -3.15561 |  |
| **KDa-pisc-s** | KDa-pisc | Tok-bn | -0.00136393 | 0.000597315 | -2.28343 |  |

Note: se – standard error; Focal – Focal population in which admixture is detected; Ref – Reference populations 1 and 2.

**Supplementary Table S7.**
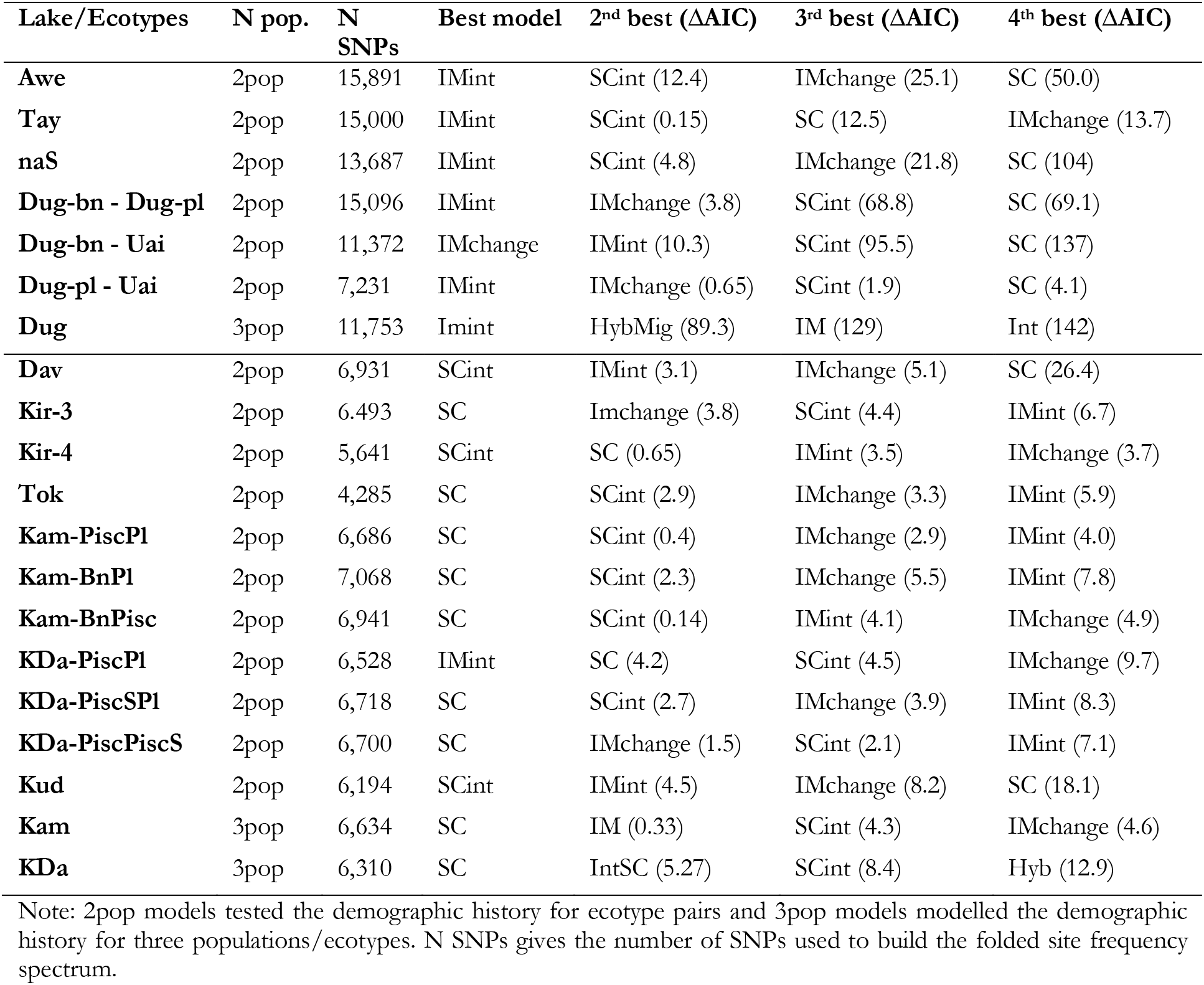
The most likely demographic models for each lake population. Shown are the fours best fitting models for each population/comparison and the respective ΔAIC between them.

**Supplementary Table S8.** Correlations of demographic parameters and trajectory lengths.

| Dem. parameter | Trajectory length | R<sup>2</sup> | P-value |
| --- | --- | --- | --- |
| **T<sub>DIV</sub>** | L<sub>P</sub> | 0.055 | 0.505 |
| **T<sub>DIV</sub>** | L<sub>Gn</sub> | 0.199 | 0.071 |
| **T<sub>DIV</sub>** | L<sub>Go</sub> | 0.334 | 0.023 |
| **T<sub>DIV</sub>** | L<sub>RDA</sub> | 0.015 | 0.305 |
| **T<sub>DIV</sub>** | L<sub>GEx</sub> | 0.372 | 0.118 |
| **T<sub>SC</sub>** | L<sub>P</sub> | 0.065 | 0.546 |
| **T<sub>SC</sub>** | L<sub>Gn</sub> | 0.045 | 0.499 |
| **T<sub>SC</sub>** | L<sub>Go</sub> | 0.026 | 0.423 |
| **T<sub>SC</sub>** | L<sub>RDA</sub> | 0.071 | 0.205 |
| **T<sub>SC</sub>** | L<sub>GEx</sub> | 0.201 | 0.709 |
| **Nm** | L<sub>P</sub> | 0.030 | 0.421 |
| **Nm** | L<sub>Gn</sub> | 0.070 | 0.195 |
| **Nm** | L<sub>Go</sub> | 0.013 | 0.305 |
| **Nm** | L<sub>RDA</sub> | 0.205 | 0.079 |
| **Nm** | L<sub>GEx</sub> | 0.186 | 0.217 |

Note: Dem. parameter – Demographic parameter.

**Supplementary Table S9.** Ecotype-associated expressed genes with associated cis-eQTL.

| Chrom | Pos | N.cis-SNPs | Gene | F-statistic | FDR | beta | R<sup>2</sup> | RDA.axis | RDA.loading |
| --- | --- | --- | --- | --- | --- | --- | --- | --- | --- |
| Contig2861 | 682439 | 1 | GENE37518_R0 | -4.3035659 | 0.07864162 | -1.1812016 | 0.39811713 | 1 | 0.082149302 |
| Contig9511 | 57765 | 9 | RNA101009_R0 | -6.9147358 | 0.00147824 | -0.669493 | 0.63067299 | 1 | -0.094215753 |
| Contig706 | 35988 | 1 | RNA124088_R1 | 4.62134923 | 0.05210489 | 1.43091656 | 0.43270307 | 1 | -0.064353881 |
| Contig2100 | 384673 | 2 | RNA13698_R0 | -4.4057006 | 0.07225572 | -0.7968371 | 0.40940977 | 2 | -0.082842073 |
| Contig12095 | 1207 | 1 | RNA14706_R0 | -4.4032784 | 0.07225572 | -0.8662297 | 0.40914385 | 2 | -0.079057491 |
| Contig587 | 1174255 | 1 | RNA17901_R0 | -4.2087891 | 0.09003047 | -0.4451682 | 0.38749491 | 2 | 0.070797588 |
| Contig1414 | 2885789 | 1 | RNA18530_R0 | 4.97030319 | 0.03375758 | 2.21531134 | 0.46873016 | 1 | 0.072009723 |
| Contig1201 | 885031 | 2 | RNA21425_R0 | -4.1555588 | 0.09419291 | -0.4453406 | 0.38147066 | 2 | 0.071708183 |
| Contig904 | 1175920 | 1 | RNA21720_R0 | -4.8734871 | 0.04020891 | -0.4691819 | 0.45894637 | 1 | 0.107174449 |
| Contig1631 | 646839 | 2 | RNA23287_R1 | 4.2395809 | 0.0875681 | 0.26035486 | 0.39096072 | 2 | 0.090023064 |
| Contig2947 | 170266 | 2 | RNA25489_R0 | -4.566751 | 0.05634307 | -0.7769089 | 0.42687797 | 1 | 0.075613456 |
| Contig1397 | 391240 | 1 | RNA26428_R1 | -4.3621683 | 0.07529024 | -0.4727803 | 0.40461651 | 1 | -0.082573446 |
| Contig1358 | 911738 | 1 | RNA27893_R0 | -4.2044756 | 0.0902104 | -0.4292859 | 0.38700828 | 1 | -0.106208673 |
| Contig699 | 787330 | 3 | RNA35235_R1 | 4.54674664 | 0.05672388 | 0.55518712 | 0.42473128 | 1 | -0.094163469 |
| Contig3135 | 697831 | 1 | RNA46270_R0 | 4.65985404 | 0.05092883 | 0.53114402 | 0.43678109 | 2 | -0.058797069 |
| Contig1700 | 1584465 | 1 | RNA54079_R0 | 4.71775654 | 0.0464547 | 1.05908525 | 0.4428662 | 1 | -0.082845396 |
| Contig3633 | 211497 | 1 | RNA62277_R0 | 5.72605989 | 0.01237167 | 1.39145353 | 0.53938097 | 2 | -0.069869143 |
| Contig4257 | 38394 | 1 | RNA65452_R1 | -5.5394866 | 0.01759705 | -3.0711062 | 0.52288379 | 2 | -0.062621388 |
| Contig603 | 1161004 | 1 | RNA71628_R0 | 4.38937387 | 0.0724093 | 0.35069496 | 0.40761556 | 1 | 0.072100135 |

Note: Chrom – Contig, Pos – SNP position, N.cis-SNPs – Number of cis-QTL associated with expression of the particular gene. FDR – false discovery rate, beta – effect size of cis-eQTL. R<sup>2</sup> – Effect size of the correlation between expression and genotype. RDA.axis – Axis of gene expression divergence from the RDA analysis (corresponds to Fig. 6d), RDA.loading – Loading of each gene along the corresponding RDA axis.

| **Chrom** | **Pos** | **N.cis-SNPs** | **Gene** | **F-statistic** | **FDR** | **beta** | **R<sup>2</sup>** | **RDA.axis** | **RDA.loading** |
| --- | --- | --- | --- | --- | --- | --- | --- | --- | --- |
| Contig1708 | 223269 | 1 | RNA81939_R0 | 4.49981188 | 0.06100233 | 0.36702121 | 0.41966876 | 1 | -0.072706216 |
| Contig698 | 1344459 | 1 | RNA83442_R0 | -4.7282695 | 0.0464547 | -0.6191383 | 0.44396489 | 2 | 0.063960238 |
| Contig1005 | 1644002 | 1 | RNA85418_R0 | 4.44869915 | 0.06718682 | 0.44987576 | 0.4141147 | 2 | 0.086576251 |
| Contig985 | 1861483 | 1 | RNA86913_R1 | -5.0103838 | 0.032617 | -1.5054908 | 0.47273221 | 1 | 0.088755726 |
| Contig1560 | 1213339 | 1 | RNA8803_R1 | -4.6744939 | 0.05026973 | -0.8396593 | 0.43832501 | 2 | -0.075225058 |
| Contig836 | 348293 | 1 | RNA97564_R0 | 4.39942336 | 0.0724093 | 0.6248348 | 0.40872044 | 2 | -0.139056588 |

Note: Chrom. – Contig, Pos – SNP position, N.cis-SNPs – Number of cis-QTL associated with expression of the particular gene. FDR – false discovery rate, beta – effect size of cis-eQTL. R<sup>2</sup> – Effect size of the correlation between expression and genotype. RDA.axis – Axis of gene expression divergence from the RDA analysis (corresponds to Fig. 6d), RDA.loading – Loading of each gene along the corresponding RDA axis.

**Supplementary Table S10.** WGCNA networks. Gene co-expression modules in network generated from 1,512 ecotype associated genes.

| Module | Number genes in module | Significantly (FDR < 0.05) associated with | Ecotype associated with |
| --- | --- | --- | --- |
| Blue | 221 | Ecotype | pl |
| Cyan | 42 | Ecotype | pl |
| Magenta | 73 | Lake + Ecotype | pl |
| MidnightBlue | 31 | Ecotype | pl |
| Salmon | 43 | lake + Ecotype | pl |
| Black | 98 | Ecotype | pl |
| Green | 159 | Ecotype | pl |
| GreenYellow | 43 | Ecotype | bn |
| LightCyan | 73 | Lake + Ecotype | bn |
| Turquoise | 384 | Ecotype | bn |
| Brown | 162 | Ecotype | bn |
| Pink | 73 | Ecotype | bn |
| Red | 110 | Ecotype | bn |

| Module | Ecotype associated with | KEGG | KEGG pathway description | FDR |
| --- | --- | --- | --- | --- |
| Blue | pl | dre00240 | Pyrimidine metabolism - *Danio rerio* (zebrafish) | 0.04 |
| Magenta | pl | dre04510 | Focal adhesion - *Danio rerio* (zebrafish) | 0.04 |
| Magenta | pl | dre04512 | ECM-receptor interaction - *Danio rerio* (zebrafish) | 0.04 |
| Black | pl | dre00561 | Glycerolipid metabolism - *Danio rerio* (zebrafish) | 0.02 |
| Black | pl | dre04350 | TGF-beta signalling pathway - *Danio rerio* (zebrafish) | 0.02 |
| Turquoise | bn | dre04070 | Phosphatidylinositol signalling system - *Danio rerio* (zebrafish) | 0.01 |
| Turquoise | bn | dre00562 | Inositol phosphate metabolism - *Danio rerio* (zebrafish) | 0.03 |

Note: Traits (lake and ecotype) significantly correlated (FDR < 0.05) with each module are given, as well the direction of expression for ecotype: pl = up-regulated in planktivorous (over benthivorous) and bn = up-regulated in benthivorous (over planktivorous). KEGG – KEGG pathway ID, FDR – False discovery rate

**Supplementary Table S11.**
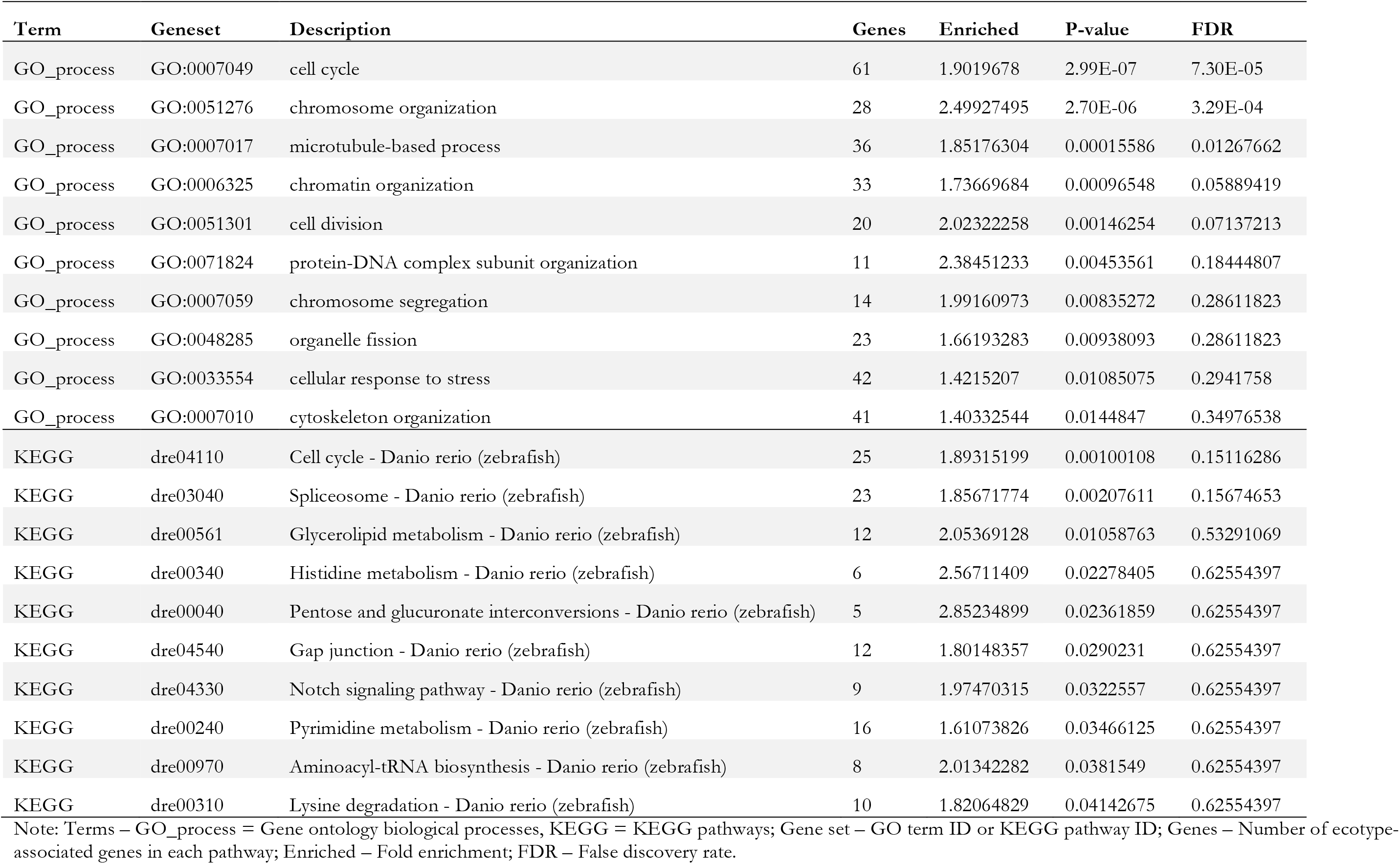
Gene ontology overrepresentation results for ecotype-associated expressed genes (RDA).

